# Trehalose mediates salinity-stress tolerance in a crustacean

**DOI:** 10.1101/2024.02.09.579427

**Authors:** Joana L. Santos, Fabienne Nick, Nikko Adhitama, Peter D. Fields, Jonathon H. Stillman, Yasuhiko Kato, Hajime Watanabe, Dieter Ebert

**Affiliations:** Department of Environmental Sciences, Zoology, University of Basel, Vesalgasse 1, 4051 Basel, Switzerland; Department of Biotechnology, Graduate School of Engineering, Osaka University. 2-1 Yamadaoka, Suita, Japan 565-0871; Institute for Open and Transdisciplinary Research Initiatives (OTRI), Osaka University, Suita 565-0871, Japan; Department of Biology, San Francisco State University. 1600 Holloway Ave., San Francisco CA 94132 USA; Department of Integrative Biology, University of California Berkeley, 3040 Valley Life Sciences Building 3140, Berkeley, CA 94720 USA

**Keywords:** Freshwater salinization, local adaptation, phenotypic plasticity, salt, stress response, trehalose, *alpha*, *alpha-trehalose-phosphate synthase*, *Daphnia magna*

## Abstract

Salinization poses an increasing problem worldwide, threatening freshwater organisms and raising questions about ability to adapt. We explore the mechanisms enabling a planktonic crustacean to tolerate elevated salinity. By gradually raising water salinity in clonal cultures from 185 *Daphnia magna* populations, we showed that salt tolerance strongly correlates with native habitat salinity, indicating local adaptation. A GWAS revealed a major effect of the *Alpha,alpha-trehalose-phosphate synthase* (*TPS*) gene, suggesting that trehalose production facilitates salinity tolerance. We found a positive correlation between water salinity and trehalose concentrations in tolerant animals, while intolerant animals failed to produce trehalose. Using CRISPR/Cas9, a silenced *TPS* gene supported the role of trehalose under salt stress. Our study highlights how a keystone freshwater animal adapts to salinity stress using an evolutionarily conserved mechanism known in plants and bacteria, but not in metabolic-active animals.

**Highlights:**

- Salinity tolerance is a locally adapted trait in *Daphnia magna*
- The trehalose *TPS* gene shows a strong association with salinity tolerance
- Trehalose content increases with water salinity allowing survival of tolerant genotypes
- Non-functional *TPS* genes reduce salinity tolerance by preventing trehalose production

## Introduction

Salinization is an emerging threat to freshwater ecosystems, causing substantial economic losses in agriculture and aquaculture^1^ and reducing biodiversity in natural freshwater habitats^2^. Since most organisms can tolerate only limited salinity ranges^4^, they are forced to adjust to local conditions by developing behavioural, physiological or structural adaptations^5^, or by migrating, when possible, to other habitats with suitable ecological conditions^6^. Failure to adapt may result in local extinction. On the other hand, adaptations to local environmental conditions may lead to genetic differentiation among populations for the genes under selection^7^.

Several molecular mechanisms allow bacteria, plants and animals to tolerate elevated salt levels^8–10^. These include transcriptional and post-transcriptional factors, including small RNAs^11^, and translational and post-translational regulators that induce protein ubiquitination^12,13^. Expression of genes that convey salt tolerance may induce physiological, biochemical and molecular responses, resulting in phenotypic alterations that manage salt uptake and transport^14,15^, regulate signalling molecules (e.g. phytohormones)^16^, influence the production and accumulation of reactive oxygen species, and modulate processes such as programmed cell death^17^. Furthermore, salinity stress can stimulate the production of proteins^18^ and solutes with osmotic and protective functions, such as proline, sucrose and trehalose^19^. As salt-induced stress is energetically demanding, carbohydrate and lipid metabolisms may also be activated^20^.

As a keystone species in standing freshwater bodies, *Daphnia magna* is a widespread crustacean across the Holarctic^21^. It has become a primary model to understand genetics in an environmental context^22,23^, by linking important ecological traits, such as resistance to environmental stressors^24–27^, to genomic regions. A powerful aspect of the *Daphnia* system is the possibility to breed animals both sexually and asexually (clonal), enabling researchers to separate genetic and non-genetic effects in common garden experiments. Here we use this system aiming to identify whether salinity tolerance is a locally adapted trait in *Daphnia* by using genotypes originating from a variety of different habitats, including estuaries, freshwater lakes and rockpools^21^ and to identify the genetic basis of salinity tolerance by performing a genome-wide-association study (GWAS). As the *Alpha,alpha-trehalose-phosphate synthase* (*TPS*) gene—involved in trehalose synthetises—showed a strong association with the studied trait, we developed experiments to detect this sugar in individuals maintained at elevated salinity, ultimately confirming, that trehalose concentration is positively correlated with salinity tolerance. Two TPS mutant lines of a single salinity-tolerant genotype developed using CRISPR/Cas9, were no longer able to produce trehalose at elevated salinity and had reduced salinity tolerance. Our findings highlight the contribution of trehalose to tolerance of salt stress and reveal how local selection has shaped this trait across the large geographic range of a keystone crustacean.

## Results

### Local adaptation of salinity tolerance

Exposing clonal lines of 185 *D. magna* populations to a gradual increase in salinity (measured as water conductivity (mS/cm)) revealed large variation in salinity tolerance among genotypes (Fig. 1, Fig. S1, Table S1).

**Figure 1.**
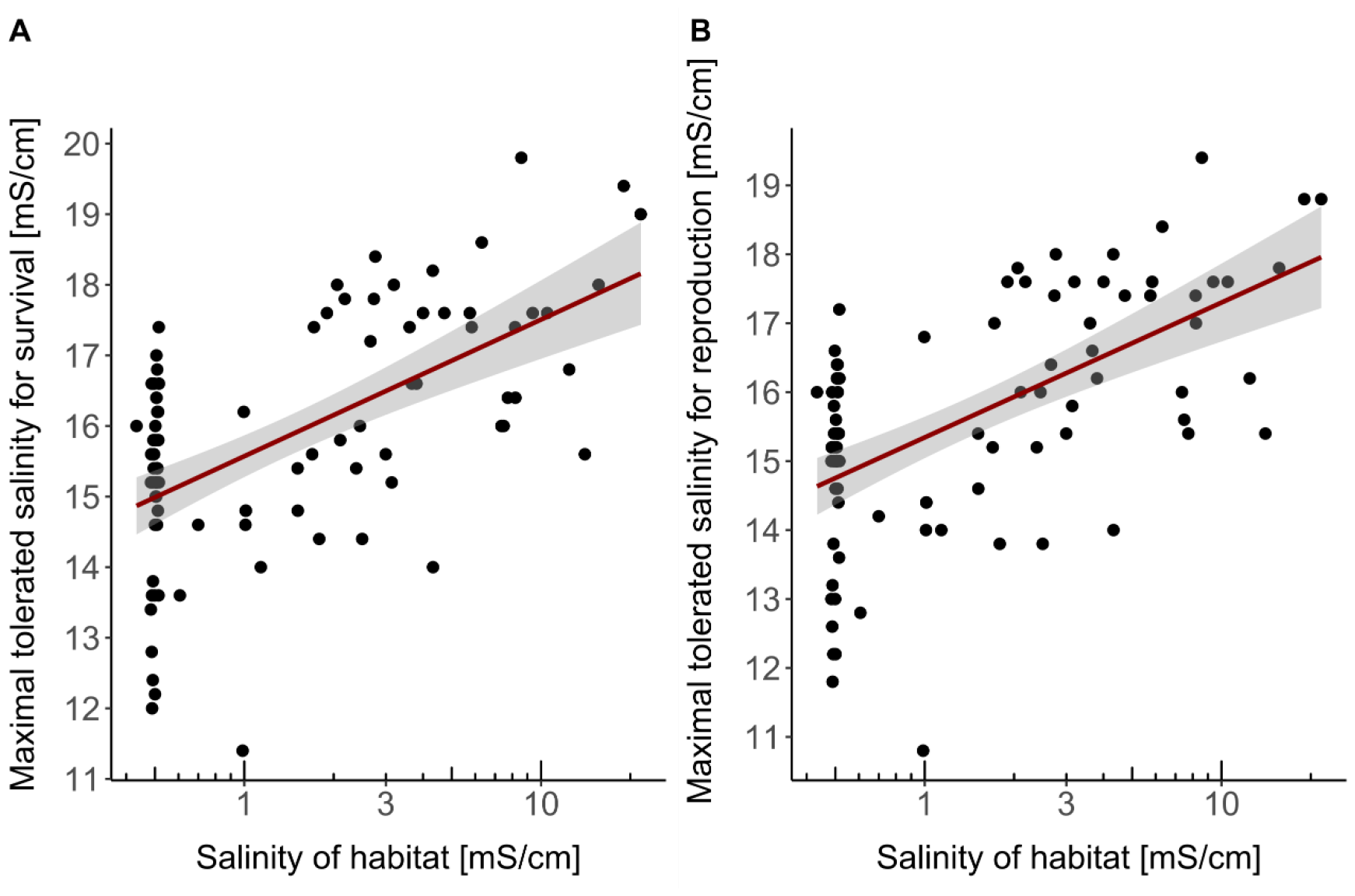
Local adaptation of salinity tolerance. Maximal recorded salinity tolerance for survival (A) and reproduction (B) in *D. magna* genotypes indicated by a correlation between tolerance and water salinity at the sampling site. The high density of samples at 0.5 mS/cm on the x-axis occurs because habitats reported as “freshwater” are assumed to have approximately this salinity level. The red line is a linear regression with its 95 % confidence interval indicated in grey shading. See also Table S1 and Fig. S1.

Maximal tolerated salinity for survival ranged from 10.0 to 19.8 mS/cm, and for reproduction (averaged for asexual and sexual reproduction) ranged from 8.4 to 19.4 mS/cm (Fig. 1, Table S1). Genotypic effects explained 87.8 % of the total variation for maximal tolerated salinity for survival (F-value=37.1, DF=768, *p-*value<2.2x10^−16^, R^2^=0.878), and 85.5 % of the total variation for maximal tolerated salinity for reproduction (F-value=30.55, Df=768, *p-*value<2.2x10^−16^, R^2^=0.855). A positive correlation was found between maximal water salinity recorded at the genotype’s site of origin and maximal tolerated salinity for survival and reproduction, confirming that genotypes from saltier habitats possess higher salinity tolerance (Fig. 1A; survival: n=90, rho=0.64, S=27507, *p-* value=4.2x10^−10^; reproduction: n=90, rho=0.64, S=27054, *p-*value=2.6x10^−10^).

### Genome-wide association study for salinity tolerance

A genome-wide association study (GWAS) revealed a major candidate gene potentially underlying salinity tolerance.

The total genomic dataset of 178 *D. magna* genotypes varied at 22,921,419 sites (32.1 % multiallelic and 67.9 % biallelic SNPs), but only 2,934,446 SNPs were used for analysis, after filtering to include only biallelic positions and to exclude SNPs with low-quality calls, with minor allele frequency lower than 0.05, single nucleotide indels and with missing data among genotypes greater than 0.9. There was a strong association between salinity tolerance and a single genomic region located on chromosome 7, contig 000003F (Fig. 2, Fig. S2, Table S2). We identified 27 SNPs within a window of 3676 base pairs above the genome-wide Bonferroni corrected significance *p-*value threshold of 3.4 x 10^−9^ (Fig. 2, Table S2).

**Figure 2.**
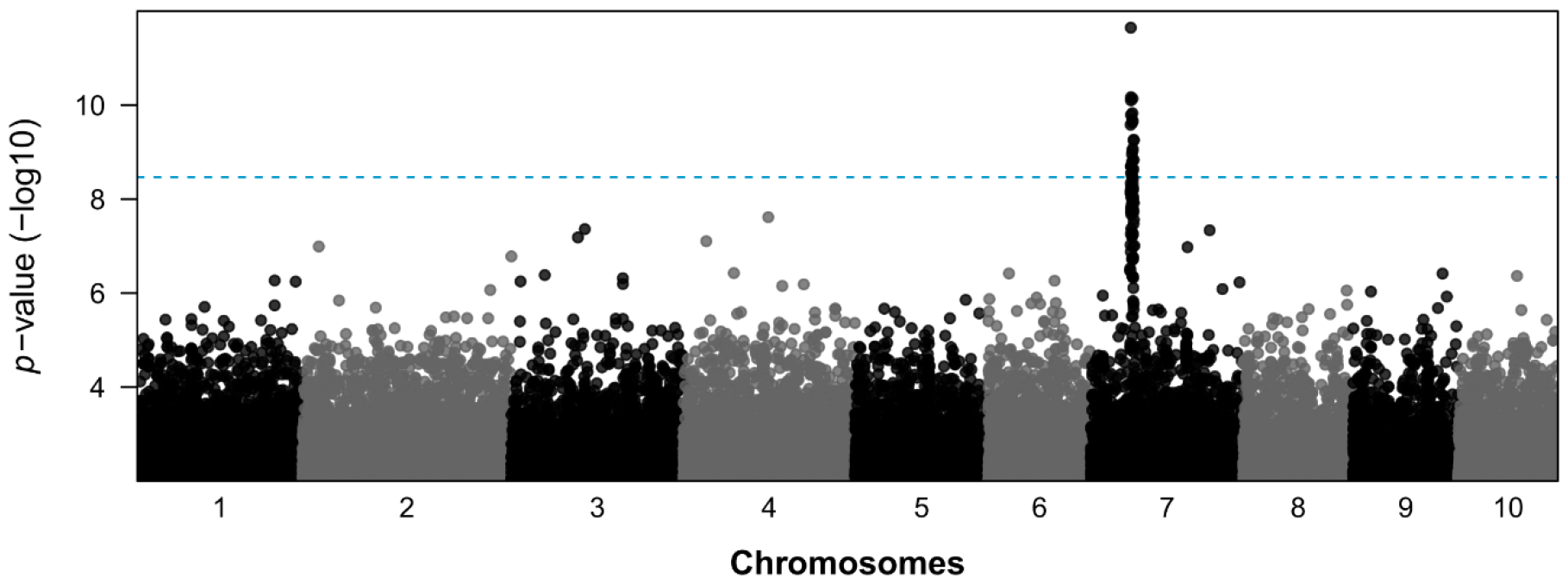
Genome-wide association study (GWAS) for salinity tolerance. The Manhattan plot shows a highly significantly associated peak region in chromosome 7, contig 000003F. The blue dashed line represents the Bonferroni adjusted *p*-value=3.4x10^−9^. The alternating pattern of black and grey dots indicates the 10 chromosomes of *D. magna*. See also Table S1, Fig. S2 and Table S4.

Using GenBank^28^ and the annotated reference genome of *D. magna* (BioProject ID PRJNA624896), we identified three genes in the peak region: the *Alpha,alpha-trehalose-phosphate synthase* (*TPS*) gene, the *guanine exchange factor Vav3* (*Vav3 GEF*) and an uncharacterized gene (GenBank accession numbers XM_032933135.1, XM_032933134.1 and XM_032933154.1, respectively) (Table S2). The two characterized genes were also identified by blasting against the EnsemblMetazoa database (http://metazoa.ensembl.org; gene number APZ42_022324 for *TPS* and APZ42_022330 for *Vav3 GEF*). The *TPS* gene was depicted in 70.4 % of the outliers SNPs (Table S2) and, since it is involved in the synthesis of trehalose, a molecule known to play a role in salt-tolerance in bacteria and plants^29,30^, we considered it as our prime candidate.

### Trehalose correlates positively with salinity levels and tolerance

While gradually increasing salinity and measuring survival, trehalose and protein concentrations in genotypes from the high and low range of salinity tolerance, we evidence trehalose production in relation to salt stress and differences among tolerant and intolerant *D. magna* genotypes (Fig. 3, Table S1, Fig. S3 and S4).

**Figure 3.**
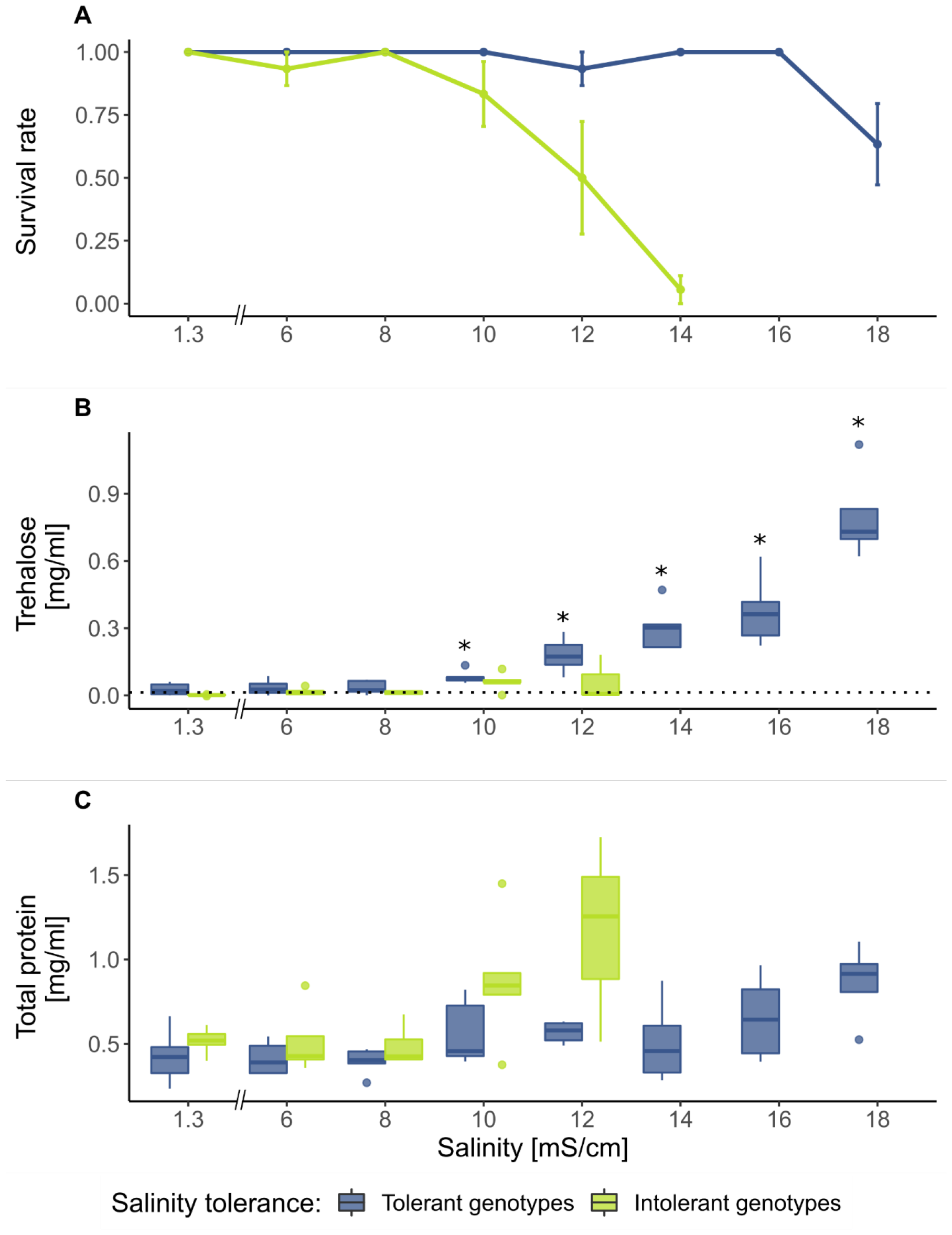
Trehalose and protein concentration relates with salinity levels and tolerance. (A) Survival rate per tolerance group and treatment, showing mean dots connected by lines and the standard error of the mean. (B) Estimated trehalose concentration per tolerance group and treatment. The dashed black line represents the mean trehalose concentration of the blank samples, and asterisks reporting significant values of trehalose being different from the blank samples for each tolerance group and treatment. (C) Estimated total dried protein concentration per tolerance group and treatment. Boxplots in (B) and (C) show the median, first and third quantile. Whiskers extend to 1.5 times the interquartile range upper and lower limits. The dots show data points beyond the whiskers. Artificial Daphnia Medium (ADaM) has a salinity of 1.3 mS/cm. See also Table S1, Fig. S2 and Fig. S3

Survival differed strongly among the genotypes chosen to represent the high and low salinity tolerance group (Fig. 3A). Intolerant genotypes died at salinity levels of 10 to 14 mS/cm, while the tolerant genotypes died at 18 mS/cm or higher. Salinity intolerant genotypes had no or insignificant 0.04±0.04 mg/ml), while salinity tolerant genotypes contained significant amounts of trehalose from 10 mS/cm salinity levels upwards (Fig. 3B; exponential regression model: overall F-statistics=7.33, *p*-value=3.8x10^−5^, Adjusted R^2^=0.54). Protein concentration also varied among treatments, showing a significant increase when salinity reached the tolerance limit (Fig. 3C; exponential regression model: overall F-statistics=3.23, *p*-value=1.8x10^−3^, Adjusted R^2^=0.31; treatment effect: Df=7, F-statistics=4.66, *p*-value=4.6x10^−4^). Salinity intolerant genotypes had a larger increase in protein concentration (Fig. 3C; tolerance group effect: Df=1, F-statistics=9.99, *p*-value=2.7x10^−3^; interaction tolerance-group vs treatment: Df=4, F-statistics=0.54, *p*-value=0.71).

### TPS mutations prevent trehalose production and reduce salinity tolerance

Two CRISP/Cas9 TPS mutant genotypes (A and B) supported the role of the TPS gene for salinity tolerance (Table S3, S4 and S5, Fig. S5).

The control and the TPS mutant genotypes differed in all variables analysed (Fig. 4). Survival of TPS mutant genotypes began decreasing at a salinity of 10 mS/cm, and all replicates went extinct at higher salinities (Fig. 4A). In contrast, the maximal tolerated salinity for the control genotypes was 16 mS/cm (Fig. 4A). Animals of the control genotypes contained significant amounts of trehalose starting at treatment 10 mS/cm, and increasing further with salinity (Fig. 4B; exponential regression model: overall F-statistics=249, *p*-value=4.8x10^−5^, Adjusted R^2^=0.99). Trehalose concentration in TPS mutant genotypes was indetectable and did not differ from the blank samples (Fig. 4B). Protein also varied among salinity treatments, as seen in the previous experiment (Fig. 4C; exponential regression model: overall F-statistics=38.15, *p*-value=4.3x10^−7^, Adjusted R^2^=0.95; treatment effect: Df=6, F-statistics=56.92, *p*-value=1.2x10^−7^), with higher protein concentrations observed in the intolerant genotypes only in their maximal tolerated salinity treatment (Fig. 4C; tolerance group effect: Df=1, F-statistics=7.55, *p*-value=0.02; interaction tolerance-group vs treatment: Df=3, F-statistics=4.94, *p*-value=0.02).

**Figure 4.**
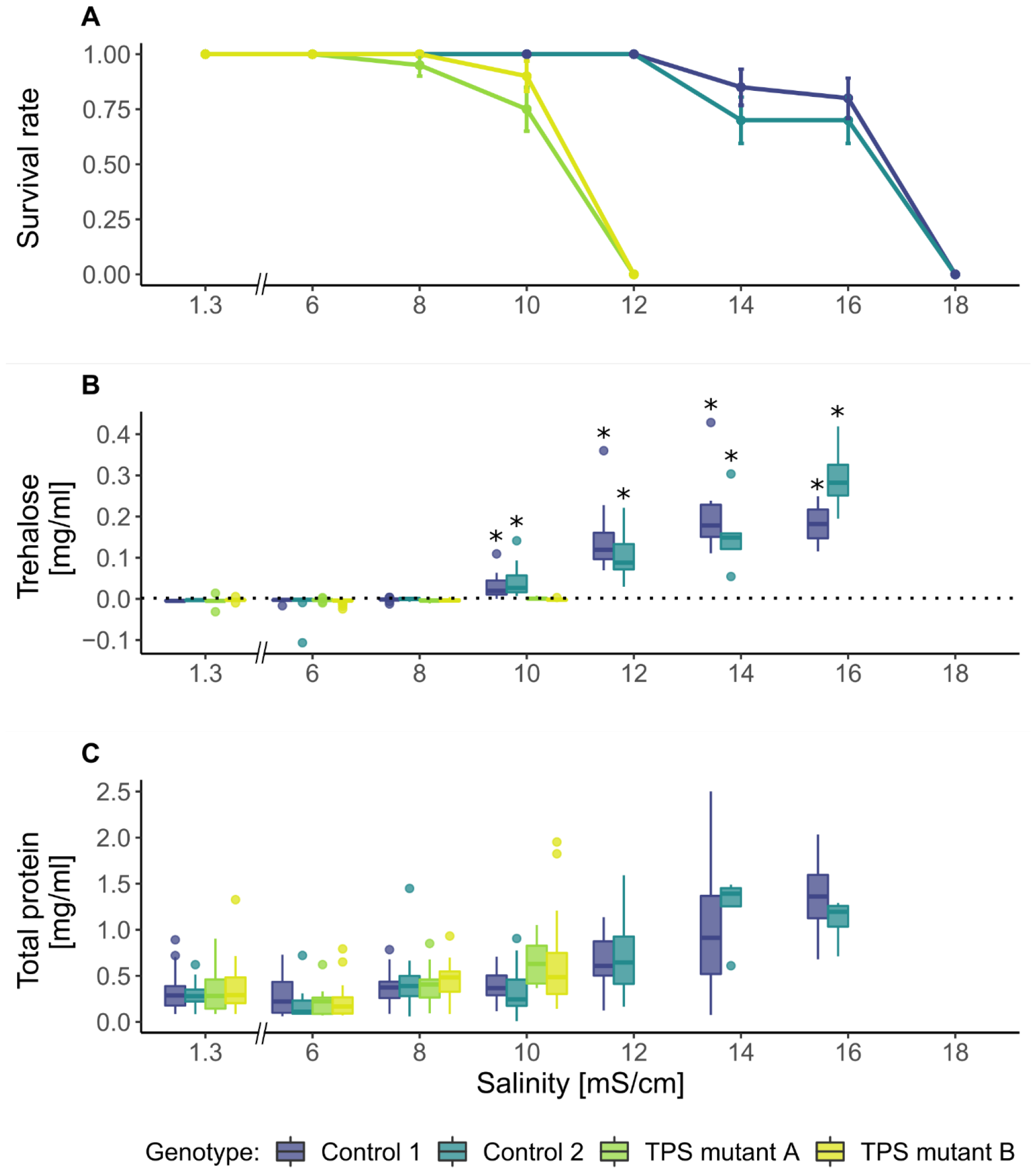
TPS mutations prevent trehalose production and reduce salinity tolerance. (A) Survival rate per genotype and treatment, showing mean dots connected by lines and the standard error of the mean. (B) Estimated trehalose concentration per genotype and treatment. The dashed black line represents the mean of trehalose concentration for blank samples and asterisks reporting significant higher values of trehalose than the blanks, for each genotype and treatment. (C) Estimated total dried protein concentration per genotype and treatment. Boxplots are as in Fig. 3. Artificial Daphnia Medium (ADaM) has a salinity of 1.3 mS/cm. See also Fig. S2, S4 and Table S2, S3 and S5.

### No link between TPS mutations and metabolic rate

The TPS mutants and control genotypes samples showed an average oxygen consumption rate of 0.35±0.01 and 0.37±0.01 µg O2/h mm^-1^, respectively. The genetic manipulation using CRISPR/Cas9 did not cause a change in routine metabolic rate, as no differences were found between TPS and control genotypes (Fig. 5; Df=1, X^2^=0.26, *p*-value=0.61) in any treatment (Fig. 5; Df=1, X^2^=0.01, *p*-value=0.94). However, salinity treatment significantly explained variation in oxygen consumption rate (Fig. 5; Df=1, X^2^=17.31, *p*-value=3.2x10^−5^), specifically, the elevated salinity treatment (conductivity 10 mS/cm) showed higher oxygen consumption rates (size corrected; mean of 0.39±0.01 µg O2/h mm^-1^) than the control treatment (conductivity 1.3 mS/cm)(mean of 0.32±0.01 µg O2/h mm^-1^). This suggests an elevated aerobic ATP production at elevated salinities. Genotypes differed significantly in their oxygen consumption rates (Df=3, *t*-value=17.1, *p*-value=3.5x10^−4^), but this concerned only the control genotypes, thus biologically irrelevant for our study (Fig. 5).

**Figure 5.**
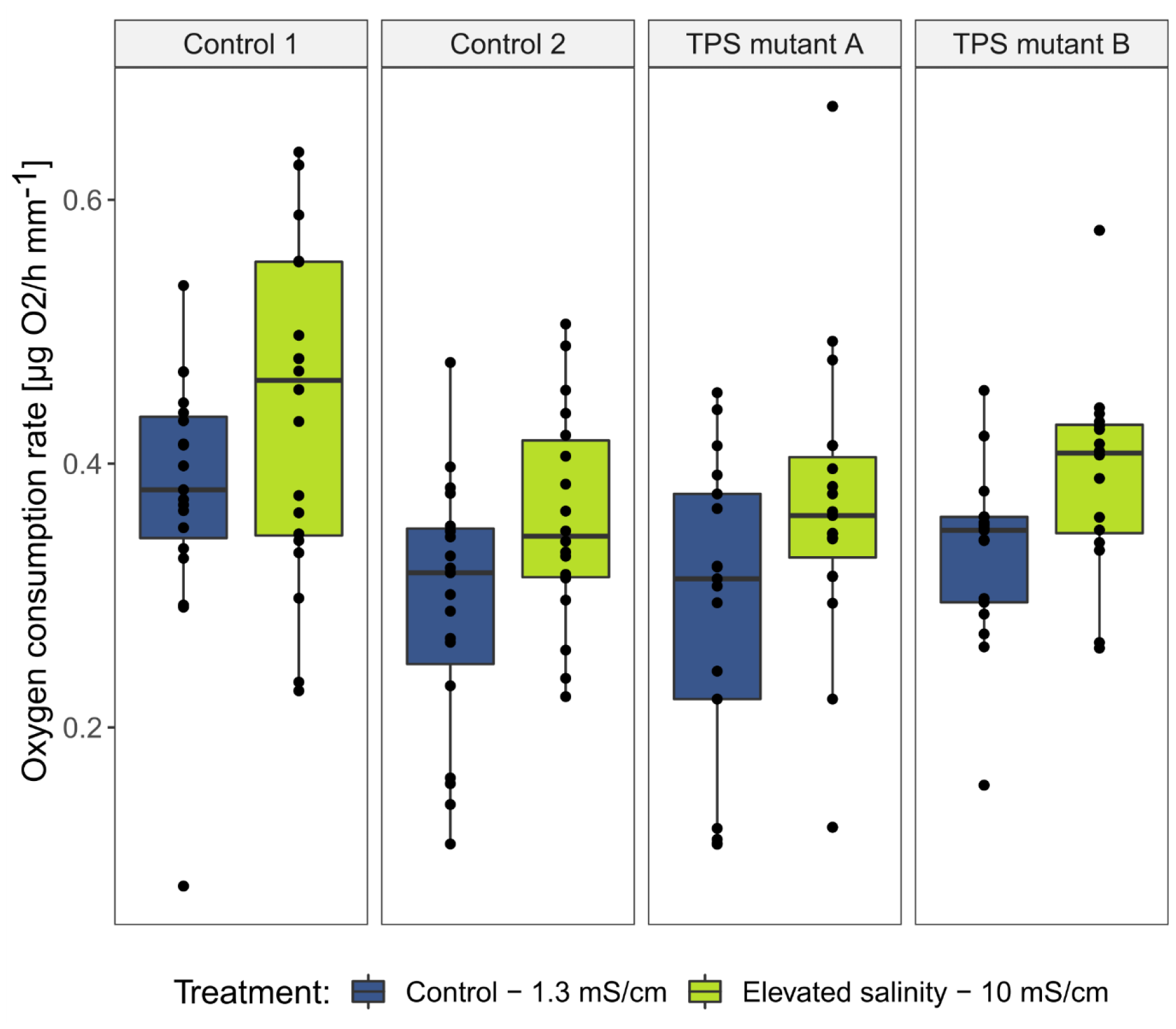
No link between TPS mutations and metabolic rate. Variation in oxygen consumption rate between control (1 and 2) and TPS mutant (A and B) genotypes at low (1.3 mS/cm) and high (10 mS/cm) salinities. Respiration rate is corrected for *D. magna* individual body length. Data are represented as mean ± SEM. Dots represent individual measurements. Boxplot are as in Fig. 3.

## Discussion

Across our planet, freshwater habitats are threatened by an increasing influx of salt, changing the conditions for many freshwater species and entire aquatic communities. Here we are asking if and how a widespread freshwater planktonic organism copes with elevated water salinities. We found large genetic variation for salinity tolerance and a clear signal of local adaptation in the genotypes we collected from 185 Holarctic *D. magna* populations, a model species widely used in ecology and evolution^21^. We found that a key player in adaptation to salinity stress is the *Alpha, alpha-trehalose-phosphate synthase* (*TPS*) gene, which is responsible for the penultimate step in the production of trehalose^31,32^, a sugar known to contribute to osmoregulation in diverse plants^30,33^, fungi^34^ and bacteria^10,29^ under salinity stress. In animals, trehalose has mainly been described for its role in desiccation resistance during dormancy and as a source of energy, but not for osmoregulation^33,35,36^.

Our study using *D. magna* genotypes from most of its Holarctic range revealed a clear correlation between salinity tolerance and the water salinity in the habitat of origin, strongly suggesting that populations are adapted to the water salinity in their local environment. Local adaptation of this model species has also been observed in response to other stressors, such as heavy metals, predators, elevated temperature, and frequency of habitat desiccation^37–40^. The strongly subdivided population structure typical for pond- and lake-dwelling species may facilitate local adaptation^21^. Our data show that a genotype’s maximal salinity tolerance has a very strong genetic component, with more than 85 % of total variation explained by genotype, highlighting the high potential of this species to adapt to changes in environmental salinity. Local adaptation to environmental salinity is also observed in other organisms including plants^41,42^, copepods^43^, molluscs^44^, fish^45,46^ and amphibians^47^, through a diversity of physiological mechanisms. Indeed, salinity stress is possibly one of the oldest environmental stress-factors on this planet.

Our GWAS uncovered a well-defined genomic region containing three genes, two of which have known biological functions. The highest number of associated SNPs fall within the *TPS* gene region. The *TPS* gene is involved in the production of trehalose, a natural sugar known for being a source of energy and especially for alleviating the adverse effects of environmental stresses such as desiccation, heat, cold and elevated salinity^35,48^. Trehalose protects and stabilizes dry proteins and membranes by filling the spaces where water molecules would have bound^49^. Trehalose also acts as a signalling molecule and elicits genes involved in detoxification and stress response^33,48^. Although trehalose is a compatible solute^31^, its biosynthesis is energetically costly^50^, and its accumulation may lead to aberrations or interfere with reactive oxygen species signalling and reducing programmed cell damage^48^. Therefore, trehalose may only be synthesized when its benefits outweigh its costs. We found that trehalose was only produced under salinity levels of 10 mS/cm or higher, and that its production increased exponentially with increased salinity, indicating that trehalose production is phenotypically plastic; it is produced on-demand and in response to an environmental cue. Plasticity in trehalose production was genotype dependent, as some genotypes produced no trehalose and were thus unable to cope with elevated salinity. This polymorphism indicates that the plasticity to produce trehalose has a cost. Specifically, animals from populations with consistently low salinity, where trehalose is never needed, may lose their ability to produce it on demand. Costs associated with plasticity can include the machinery for its upkeep or the potential misinterpretation of environmental cues^51^. The latter seems unlikely, as osmotic stress should be easy to assess with little error (unlike other cues, such as sensing and responding to potential predators). Alternatively, if the mechanism to produce trehalose is never used, the underlying genes may accumulate mutations that are neutral in the habitat where they arise, but are deleterious under conditions of elevated salinity. This hypothesis implies that salinity intolerance is a derived state and that the ancestors were salinity tolerant.

TPS is an enzyme involved in the penultimate step of trehalose production: it synthetizes trehalose-6-phosphate from UDP-glucose and glucose-6-P, which later is converted into trehalose by the *trehalose-6-phosphate phosphatase* (*TPP*)^31,32^. This pathway is unique in eukaryotes, whereas bacteria show distinct biosynthesis routes^31^. Our findings strongly confirm the role of trehalose in mediating the survival of salinity tolerant genotypes in elevated salinities. By inhibiting the expression of the regular *TPS* gene, we showed that mutant genotypes could neither produce trehalose nor survive under elevated salinities. Oxygen consumption rates also showed that the TPS mutant genotypes had similar routine metabolic rates as the control genotypes, indicating that differences in trehalose content and salinity tolerance between mutants and controls was due to the absence of the functional *TPS* gene rather than non-specific energetic shifts. Several studies have confirmed the role of both *TPS* and *TPP* genes in regulating trehalose production under stress response in eukaryotes^52–54^, which is likely similar to *D. magna*. Elevated environmental salt concentrations are known to trigger trehalose production in a diversity of organisms, including bacteria, plants, fungi and animals^34,48,55^. When plants are treated with exogenous trehalose or genetically modified to accumulate higher trehalose levels, their growth and photosynthesis parameters are boosted under elevated salt concentrations^56,57^. When mycorrhiza fungi produce trehalose in response to higher salinities, the salinity tolerance of the plant symbiont has also been shown to increase^58^. The link between trehalose and water salinity in animals is less clear. Although some studies have connected trehalose or trehalose-related genes, such as the genes involved in trehalose production or its transport, with altered salinity^9,59^, others have failed to do so^60–62^. This discrepancy may arise because, in animals, trehalose plays an important role in protecting resting life stages against heat, freezing and anhydrobiosis^35,36,55,63^. Since the stress from increased salinity often induces the production of resting life stages, it is not always clear whether trehalose-related genes are expressed to regulate the osmolarity of body fluids or to equip the resting life stages with trehalose in preparation for diapause. Our study allowed these two hypotheses to be uncoupled. Under salinity stress, *D. magna* rarely produce resting eggs, and the NIES genotype used in our genetic manipulation experiment never produced resting stages in our cultures or during the experiments. Thus, we believe that the trehalose production in *D. magna* was indeed linked to the osmotic stress caused by salt in the water.

Our analysis also unveiled a substantial increase in protein content closer to the maximal tolerated salinity in all *D. magna* genotypes. In intolerant genotypes this occurs at lower salinity levels. Increase in protein content may also be an adaptive response to environmental salinity, only triggered when the animals experience an osmotic stress.

Generally, routine metabolic rate decreases under physiological stress as a defense mechanism to reduce oxidative stress^64^, which was previously reported in other *Daphnia* species exposed to elevated salinity^65,66^. However, here we denoted an increase in the elevated salinity treatment for all genotypes. This increase might be explained by the need to allocate more energy to hyperosmoregulation through costly physiological processes, for example by the synthesis of trehalose and other metabolites, to overcome salinity stress (*see* Cochran *et al*.^67^).

The second gene with biological functions identified by our GWAS—the *Vav3 GEF* gene—is in part localized in the same genomic region and overlaps with the *TPS* gene region. For instance, both coding regions are annotated in the *D. magna* genome with approximately a 1000 bp distance, and the 5’ UTR region of *Vav3 GEF* overlaps with the coding region of the *TPS* gene. The *Vav3 GEF* gene is part of the Rho GTPases family, which is involved in signalling and cytoskeletal pathways^68^, mainly by participating in and coordinating cellular responses to extracellular stimuli. It serves as a key regulator of both endothelial barrier and genomic stability^69,70^, and has thus been associated with oxidative stress responses and the development of pathophysiological disorders, such as human cancer^71^. Despite the suggested role of certain guanine exchange factors with salinity stress^72^, to our knowledge, no published reports associate the specific *Vav3 GEF* gene with salinity tolerance. Therefore, we posit that the *Vav3 GEF* might have been identified in the GWAS study not necessarily due to its direct influence on salinity tolerance but rather because of its proximity to the *TPS* gene.

## Conclusion

Our findings highlight the role of trehalose in coping with salinity stress in an aquatic invertebrate. When the *TPS* gene was rendered non-functional, the mutant genotypes were left without trehalose production and where unable to respond plastically, resulting in drastically decreased salinity tolerance. In the *Daphnia* model salinity tolerance is shaped by local adaptation and phenotypic plasticity, meaning that only salinity tolerant genotypes are able to produce trehalose, but its expression is dependent on the adaptation to the salinity of the environment. Understanding how freshwater organism adapts to increasing salinity using this evolutionarily conserved mechanism will inspire further research on salinity tolerance in other animal species, helping us to understand the potential for adaptation and limits of it in a changing world. This would also benefit the aquaculture industry and help predict the fate of freshwater ecosystems that face growing salinity threats.

Acknowledgments

We thank Jürgen Höttinger, Heidi Schiffer, Urs Stiefel, Michelle Krebs and Alix Thivolle for the technical support during experimental work. This work was supported by a grant from the Swiss National Science Foundation to DE (grant number: 310030_188887).

## Author’s contributions

FN and DE designed and FN performed the salinity tolerance experiment; FN, JLS and PDF performed the GWAS study; JLS and DE designed and JLS performed the trehalose and protein quantification experiments; NA, KY and WH developed the TPS mutant genotypes; JLS and JHS designed and performed the oxygen consumption rate experiment. FN, JLS and NA wrote the respective methods and results section. JLS and DE wrote the first version of the manuscript and all authors commented on it.

## Methods

### Study system

*Daphnia magna*, Straus 1820, is a small crustacean that inhabits permanent and intermittent rockpools, rain ponds, estuaries and lakes of the Holarctic^21,73^. Previous research has indicated that the global distribution of *D. magna* is highly structured across North America, Europe, North Africa and Asia^74,75^. *Daphnia magna* has a short generation time of 8-15 days, so populations can increase rapidly by parthenogenetic reproduction and switch, under unfavourable environmental conditions, to sexual reproduction^76^.

### Salinity tolerance among D. magna genotypes

This experiment tested the salinity tolerance of small *D. magna* populations—i.e., their ability to reproduce and survive in water with elevated salt concentrations. We selected 185 genotypes from distinct Holarctic populations of *D. magna*, including a range of natural habitats with different salinity levels (Table S1). Salinity data, measured as water conductivity (mS/cm), is available for 90 of the 185 sites of origin of these genotypes, either measured directly during sampling or from published studies. Waterbodies reported as “freshwater” by the collector were assumed to have a salinity level of 0.5 mS/cm. Salinity measurements in this dataset range from 0.438 to 21.7 mS/cm.

Animals were exposed to increased salinity. Every two-weeks, ten individuals—neonates and gravid females—were transferred to a new jar with medium adjusted salinity 1 mS/cm higher than the previous one (ramp experiment; Fig. S1). If there were less than ten individuals present, all individuals were transferred. The two-week intervals represent approximately one asexual generation, enough time for animals to acclimatize to the higher salinity.

*Daphnia* artificial medium, ADaM, has a salinity of 1.33 mS/cm^77^. Salinity was increased by adding a concentrated sea salt solution (100 g salt/L) (hw-Marinemix reefer®, hw-Wiegandt GmbH, Germany), while measuring water conductivity with a SevenMulti conductivity meter (Metter Toledo). At the start of the experiment, all genotypes were simultaneously pre-acclimated for two weeks to a salinity level of 6 mS/cm. At two weeks, they were transferred to 8 mS/cm, and then, after another two weeks, to 10 mS/cm. After that, salinity was increased by 1 mS/cm every two weeks (*see the experimental design in* Fig. S1). During the experiment, replicate populations were kept in 360-mL jars, grouped in trays of 12 jars each, in a walk-in chamber with a 16h day:8h night cycle at 20 °C and 80 % of humidity. To reduce evaporation, trays were covered with transparent plastic sheets. Sporadic measurements revealed a salinity increase of about 2% in the jars over the two-week period, which was considered negligible and affected all replicates equally. Tray positions were shifted three times a week to reduce position effects. Animals were fed three times a week with a suspension of 50 million cells of *Tetradesmus obliquus* green algae per jar. When less than six animals remained in the jar, the food was reduced proportionally to avoid overfeeding. A population (replicate) was considered extinct when no animals showed any heartbeat or movement.

For each of the 185 genotypes, five replicate populations for a total of 925 populations were used. Every two weeks, the number of transferred animals, the number of females carrying eggs, and the presence of resting eggs in each replicate were recorded. Extinct replicates were noted. The end of the experiment was marked when the last surviving replicate population went extinct at 22 mS/cm. The maximal tolerated salinity level for survival was estimated as the highest level at which at least one animal of a replicate was able to survive the two-week period (i.e., salinity tolerance). Likewise, maximal tolerated salinity level for reproduction was defined as the highest salinity in which reproduction (sexual or asexual) was observed, identified as the production of sexual resting eggs and the presence of gravid females with asexual eggs or newborn animals at the end of the two-week period.

All statistical analyses were performed using the R software version 4.0.2^78^ and RStudio v. 1.3.1073^79^. The maximal tolerated salinity for survival and reproduction for each replicate were used as dependent variables. The genetic variance was calculated with the *lmer* function of the R package nlme v.3.1-148^80^ using genotype as a random effect. Residuals were tested for normality. To assess the relation between the two dependent variables and salinity at the genotype’s site of origin, the mean tolerance was calculated across all five replicates of the 70 genotypes with habitat salinity data available. The resulting genotype estimates were correlated with the water salinity at the genotype’s site of origin using the Spearman test. Data visualization was generated with the R package ggplot2 v.3.4.4^81^. Mean values are presented with the standard error of the mean, unless otherwise stated.

### Genome-wide association study for salinity tolerance

*As D. magna* genotypes expressed major differences in salinity tolerance, we conducted a genome-wide association study (GWAS) using maximal tolerated salinity for survival as the dependent variable to uncover the genetic architecture underlying salinity tolerance.

Whole genome-sequencing data was used from 178 *D. magna* genotypes (Table S1), obtained through Illumina-based whole genome sequencing. The methods for generating individual Illumina DNA libraries, DNA sequencing, and quality assessment are described in Fields *et al*.^82^. Sequence reads were mapped to the v.3.0 *D. magna* reference genome (BioProject ID PRJNA624896). The variant call format (VCF) file using GATK v.3.8^83^ ‘HaplotypeCaller’ according to GATK Best Practices recommendations^84,85^. Using VCFtools v.0.1.16^86^, the set of variant calls was filtered to include only biallelic SNPs. Additionally, SNPs with low-quality scores (DP<3, GQ<25), with minor allele frequency below 0.05, single nucleotide indels were excluded.

PLINK 2.0^87^ was used for the GWAS. Briefly, the filtered VCF file was converted to PLINK2-specific format pgen, pvar and psam files. Further filtering was applied to exclude variants with missing data above 90%, using --geno flag. The evaluation of individual relatedness among samples showed only few samples highly related among them, which can be explained by their geographic proximity. The results did not change when excluding those samples, thus analysis were performed with all samples. Since *D. magna* populations are highly structured, and show a signature of isolation by distance^74,75,88^, analysis controlled for population structure, by performing a Principal Component Analysis (PCA) using PLINK2. Our GWAS implemented the first ten Principal components in a linear model, using --glm flag. Association summaries were accessed in the form of Manhattan and Quantile-quantile (QQ) plots using BoutrosLab.plotting.general v7.0.3^89^ and qqman v0.1.9^90^ R packages, respectively. Outliers SNPs were identified applying a Bonferroni-adjusted *p*-value associated to 0.01 threshold. For an evaluation of the *p*-values distribution—in the QQ plot—a less stringent threshold to incorporate the entire peak region was applied.

Lastly, to determine the underlying genetic basis, the potential candidate genes within the significant SNP region were identified by extracting the DNA sequence and using homology searches against BLAST^91^ with default parameters to query the nr/nt database and the EnsemblMetazoa genomic sequence database^92^.

### Trehalose and protein quantification in different salinity treatments

The GWAS showed a strong association between salinity tolerance and the TPS gene, which is involved in the penultimate step of trehalose biosynthesis^31^, suggesting that *D. magna* may produce trehalose in relation to salinity stress. We thus aimed to quantify the amount of trehalose in salinity-tolerant and intolerant *D. magna* genotypes under conditions of elevated salinity, hypothesizing that the amount of trehalose present in adult animals correlates with water salinity, and that salinity-tolerant and intolerant *D. magna* genotypes differ in this respect.

Ten *D. magna* genotypes from the previous experiment were selected—five showing low estimates of maximal tolerated salinity for survival (10 – 12 mS/cm) and five showing high (around 18 mS/cm, *see* Table S1). Using six replicates per genotype, the protocol described above was followed, except that salinity was increased in steps of 2 mS/cm every two weeks (*see the experimental design in* Fig. S3).

Furthermore, with every salinity level increase, replicates were split, and one line was kept at the same salinity while the other line (11 animals) was placed in a medium at the next higher salinity level. Therefore, the number of replicate populations increased every two weeks when the next higher salinity treatment was added.

After sixteen weeks, all populations experienced at least two weeks in each salinity level unless they had gone extinct earlier or had not produced offspring. At this time, eleven females in the late juvenile or early adult stage were collected from each surviving replicate population and placed into a 1.5-mL Eppendorf tube. Pilot experiments had shown that eleven animals were sufficient for colorimetric quantification of both trehalose and protein. For each sample of eleven females, water was removed and samples were dried room temperature for about 3 hours until no water drops were visible in the tube. Then, 50 µl of ultra-purified water was added, and animals were homogenized with a small pestle and frozen for further processing within a month’s time. To understand potential variation in trehalose over time, animals from the 1.3 and 12 mS/cm treatments were collected every two weeks. This time series experiment did not reveal significant variation in trehalose content over time and will not be further discussed (Fig. S4).

For each sample, trehalose was extracted and quantified using a Megazyme trehalose kit (Megazyme, Bray, Ireland) according to Santos and Ebert^36^, and protein content was quantified using the Pierce™ Coomassie (Bradford) Protein Assay Kit (ThermoFisher Scientific, Switzerland).

Survival, trehalose and protein content were recorded for each sample at each salinity level and averaged these traits per genotype and treatment. Averages of the salinity tolerance group factor (tolerant or intolerant) were compared for each salinity level. Non-parametric Mann-Whitney U-tests were conducted to determine which samples showed significantly elevated concentrations of trehalose as compared to the blank samples. To test for increasing trehalose and protein with increasing salinity, exponential regression models using the *lm* function were fitted for trehalose and protein concentration considering salinity treatment, tolerance group and their interaction. The significance of the factors was assessed using *Anova* function of car v3.0-11^93^ package. Data analysis and graphics used additional R packages cowplot v1.1.1^94^, data.table v1.14.2^95^ and forcats v0.5.1^96^.

### TPS gene mutations using CRISPR/Cas9

To confirm the TPS gene’s role in trehalose content and salinity tolerance in *D. magna* adults, we produced mutant genotypes for this gene. For this, we used a *D. magna* genotype originated at the National Institute of Environmental Studies (NIES, Tsukuba, Japan), previously identified as tolerant to elevated salinity, with a maximal tolerated salinity at about 15.5 – 16 mS/cm. This genotype has been widely used in molecular studies and incorporates a genetic modification to express the green fluorescent protein^97^.

TPS mutant genotypes were generated by clustered, regularly interspaced, short palindromic repeats (CRISPR/Cas9) as described by Nakanishi *et al*.^98^. Two independent single-guide RNA (sgRNA) target sites were designed to target the catalytic regions of the TPS gene (Fig. S4A). These were constructed by synthesizing synthetic oligonucleotides containing a T7 promoter sequence, a gene-specific target sequence, and the first 20 nucleotides of the Cas9 binding scaffold (Table S2). The sgRNAs were then in vitro transcribed using the cloning-free method as described by Mohamad Ishak *et al*.^99^. Each sgRNA (2µM) and Cas9 protein (1µM) was mixed, incubated at 37 °C for 5 minutes to allow the formation of ribonucleoprotein complex, and injected into asexual female eggs following established procedures^97,100^. Genomic DNA was isolated, amplified using the primers in Table S3, and the presence of indels was confirmed using native PAGE gel electrophoresis. From there, the PCR products were cloned using Zero Blunt TOPO PCR Cloning Kit (Invitrogen) and sequenced. Out of fifteen successfully injected eggs, eight hatched; from those, seven developed into adults, and six produced offspring. Of those six G0 founder lines, five produced offspring with a mutation in sgRNA#2 target sites. Of these, two mutant candidates showing the clearest indel pattern were selected. The TPS A mutant genotype is a monoallelic mutant with a 2 bp frameshift deletion in the first allele and a 45 bp non-frameshift deletion in the second allele (Fig. S5B). The TPS B mutant genotype is also monoallelic, with a larger frameshift deletion (51 bp) in one allele and 27 bp non-frameshift insertion in the second allele (Fig. S5B). In both TPS mutant genotypes, alleles are predicted to produce a TPS protein with an altered glucose-6-phosphate (G6P) entry site (Fig. S5C, Table S5).

### Salinity tolerance, trehalose and protein quantification in TPS mutant genotypes

To understand the role of the TPS gene on salinity tolerance and on trehalose and protein content, we conducted an experiment using the above protocols, with the TPS mutant genotypes, A and B, versus two control genotypes (i.e., wild type genotypes for the TPS gene). The control genotypes are identical genetic lines, but the control 1 is the one from which TPS mutant genotypes were derived and has been in the laboratory of NIES, Japan, while control 2 has been kept separately for about six years at the University of Basel, Switzerland.

Per genotype, 20 replicates were selected, resulting in 80 replicate populations. Salinity level of the culture medium was increased every two weeks as described above (*see* Fig. S3). Population extinction was checked at regular intervals and samples were collected two weeks after the maximal tolerated salinity for survival was reached for the last replicate population(s). Data were analysed for population survival and for trehalose and protein content in populations across salinity treatments and compared across the four genotype lines (2 controls and 2 mutant genotypes) using a statistical approach similar to the previous experiment.

### Quantification of metabolic rate in mutant genotypes

Routine metabolic rate—estimated from the oxygen consumption rate—can shift as a consequence of organismal response to stress^64^. We determined if the TPS gene mutations influenced the metabolism of *D. magna* individuals by comparing the oxygen consumption rate of the mutant genotypes (TPS A and B) and the two control genotypes raised in the previous experiment at salinities of 1.3 and 10 S/cm. Twenty adult females per genotype and treatment were isolated into individual 100-mL jars. From those, an adult female from the second clutch was selected about five days before the oxygen consumption essay. In total the experiment comprised 160 animals (4 genotypes x 2 treatments x 20 replicates).

The oxygen consumption rate (µg O_2_/h) was calculated based on the oxygen decrease within the vials using the SDR device (PreSens, Germany) and 1-ml vials with 3 mm optode oxygen sensor spots. Oxygen consumption was measured at 1.3 and 10 mS/cm treatments simultaneously, for a total of four measurement blocks each with 24 samples per salinity treatment. We recorded percentage of oxygen saturation every 15 seconds for 45 minutes at 20 °C in each single-animal vial (or empty vial for the blanks) using the PreSens SDR software. The first 10 min of each recording were discarded from measurement. The mean of blanks (only ADaM, without animals) per plate. In total, the oxygen consumption rates of 145 animals were used, since twelve animals and one blank showed strong and unexplained abnormalities or an unfitted linear regression model with R^2^ values below 0.85 (the mean of R^2^ value is 0.96±9.5x10^−3^). Furthermore, three animals were lost during handling. Oxygen consumption rates were transformed from % O_2_/h to µg O_2_/h by using saturations of 9.05 and 8.79 mg/L O_2_ at 100 % oxygen saturation of 1.3 and 10 mS/cm salinity water at 1 atm atmospheric pressure, respectively, and considering the respirometer vials volume of 1 mL. These values were corrected for individual animal length.

For data analysis and visualization, averaged values were used per sample and grouped by genotype and salinity treatment. The effect of salinity treatment, genotype-type (TPS mutant or control), and genotype (considered as a random effect nested in genotype-type) on oxygen consumption rate (µg O2/h mm^-1^), was examined using the *lmer* function of lme4 v1.1-23^101^. The significance of fixed and random effects was accessed using *Anova* function of car v3.0-11^93^ package and afex v1.3-0^102^ package. Additional R packages were used for these analyses: tidyverse v1.3.1^103^, plyr v1.8.6^104^, readxl v1.3.1^105^, nls.multsart v1.2.0^106^ and broom v0.7.6^107^.

