## Supplementary Material for "Trehalose mediates salinity-stress tolerance in a crustacean"

ORCIDs:
0000-0003-2939-7091 (JLS); 0000-0002-5611-4387 (FN); 0000-0001-7864-7654 (NA); 0000-0003-2959-2524 (PDF); 0000-0002-4783-3830 (JHS); 0000-0002-1041-1044 (YK); 0000-0001-9657-6554 (HW); 0000-0003-2653-3772 (DE)

Table S1. *Daphnia magna* genotypes used in this study. This includes GPS location, country of origin and habitat salinity (mS/cm) when known (otherwise stated as "-"). The maximal salinity tolerated for survival (MSS) and reproduction (MSR) measured in the first experiment is provided in mS/cm. Genotypes with genome available and included in the GWAS are assigned to a "Y", otherwise with a "N", in the respective column. Genotypes with • were used in the trehalose quantification experiment.

| Genotype | Latitude | Longitude | Country | Habitat salinity | MSS | MSR | GWAS |
|---|---|---|---|---|---|---|---|
| AM-AR-1 | 39.752 | 44.806 | Armenia | - | 16.0 | 15.8 | Y |
| BE-HO-1 | 50.145 | 5.077 | Belgium | - | 15.2 | 14.6 | Y |
| BE-OHZ-T10 | 50.839 | 4.655 | Belgium | - | 15.8 | 15.8 | Y |
| BE-OM-1 | 50.863 | 4.721 | Belgium | - | 16.0 | 16.2 | Y |
| BE-T1-2 | 50.823 | 4.594 | Belgium | - | 12.2 | 11.8 | Y |
| BE-WE-G59 | 51.068 | 3.774 | Belgium | - | 16.2 | 16.2 | Y |
| BE-WH1-2 | 51.336 | 3.349 | Belgium | - | 18.0 | 17.6 | Y |
| BE-WH2-11 | 51.335 | 3.348 | Belgium | - | 17.0 | 17.0 | Y |
| BE-WTE-1-1 | 50.828 | 4.639 | Belgium | - | 17.8 | 17.8 | Y |
| BY-G-9 | 52.421 | 31.014 | Belarus | 0.500 | 14.6 | 14.6 | Y |
| CA-CAL1-1 | 50.931 | -113.865 | Canada | - | 15.0 | 14.6 | Y |
| CA-CAL2-1 | 50.937 | -113.869 | Canada | - | 14.8 | 14.4 | Y |
| CA-CH-1 | 58.771 | -93.851 | Canada | - | 13.4 | 12.6 | Y |
| CA-LL1-1 | 52.409 | -105.006 | Canada | - | 12.6 | 12.6 | Y |
| CA-LL2-2 | 52.457 | -105.049 | Canada | - | 14.2 | 13.6 | Y |
| CA-RI-1 | 48.445 | -68.585 | Canada | - | 15.0 | 15.0 | Y |
| CA-SS-1 | 51.560 | -106.357 | Canada | - | 14.4 | 14.2 | Y |
| CH-H-1 | 47.558 | 8.863 | Switzerland | 0.438 | 16.0 | 16.0 | Y |
| CN-W1-1 | 30.562 | 114.390 | China | 0.500 | 16.6 | 16.0 | Y |
| CY-PA1-1 | 35.031 | 33.974 | Cyprus | 3.680 | 16.6 | 16.6 | Y |
| CY-PA2-1 | 35.033 | 33.955 | Cyprus | - | 16.4 | 16.6 | Y |
| CY-PA3-1 | 35.034 | 33.955 | Cyprus | - | 16.0 | 15.6 | Y |
| CZ-DO-1 | 49.006 | 14.441 | Czech Republic | - | 16.2 | 16.2 | Y |
| CZ-KO-1 | 50.125 | 14.869 | Czech Republic | - | 15.6 | 15.4 | Y |
| CZ-LE1-1 | 48.783 | 16.790 | Czech Republic | - | 17.0 | 16.4 | Y |
| CZ-N1-1 | 48.775 | 16.724 | Czech Republic | - | 15.8 | 15.4 | Y |
| CZ-N2-6 | 48.767 | 16.742 | Czech Republic | - | 16.2 | 16.0 | Y |
| CZ-NVR-1 | 49.007 | 14.442 | Czech Republic | - | 16.2 | 16.4 | Y |
| CZ-SVR-1 | 49.011 | 14.437 | Czech Republic | - | 16.2 | 16.0 | Y |
| DE-G1-106 | 54.282 | 10.967 | Germany | - | 16.8 | 16.8 | Y |
| DE-GB-1 | 54.329 | 10.627 | Germany | - | 17.0 | 17.0 | Y |
| DE-K35-Mu10 | 48.207 | 11.710 | Germany | 0.500 | 13.6 | 13.6 | Y |
| DE-KA-F28 | 50.935 | 6.928 | Germany | - | 16.0 | 15.8 | Y |
| DE-KN1-4 | 54.177 | 10.807 | Germany | 0.500 | 15.4 | 15.4 | Y |
| DE-R1-1 | 54.208 | 10.424 | Germany | 0.500 | 15.2 | 15.0 | Y |
| DE-S1-8• | 48.780 | 9.183 | Germany | 0.500 | 12.8 | 12.6 | Y |
| DE-S2-1 | 48.806 | 9.173 | Germany | 0.500 | 16.4 | 16.4 | Y |
| DE-S3-3 | 48.806 | 9.172 | Germany | 0.500 | 15.6 | 15.2 | Y |
| DE-SPO-1 | 54.286 | 8.650 | Germany | - | 18.0 | 17.6 | Y |
| DK-RL-3 | 55.964 | 9.596 | Denmark | 0.500 | 15.0 | 15.0 | Y |
| DZ-JV-2 | 36.880 | 7.757 | Algeria | - | 18.0 | 17.8 | Y |
| EG-ELIAS-1 | 28.544 | 33.974 | Egypt | - | 16.4 | 16.2 | Y |

| | | | | | | | |
|---|---|---|---|---|---|---|---|
| ES-D-BDE1-F9 | 37.148 | -6.037 | Spain | - | 17.8 | 17.6 | Y |
| ES-D-CC-NF7 | 37.071 | -6.272 | Spain | - | 16.0 | 15.6 | Y |
| ES-D-H-F3 | 36.872 | -5.862 | Spain | - | 18.0 | 17.6 | Y |
| ES-HT-1 | 38.775 | -1.410 | Spain | 2.760 | 18.4 | 18.0 | Y |
| ES-HY-1 | 38.779 | -1.435 | Spain | 4.305 | 18.2 | 18.0 | Y |
| ES-OM-1[•] | 37.254 | -6.969 | Spain | 6.310 | 18.6 | 18.4 | Y |
| ES-RO-1 | 37.128 | -6.482 | Spain | - | 18.6 | 18.2 | Y |
| ES-RV-1 | 40.659 | 0.775 | Spain | - | 18.2 | 18.2 | Y |
| FI-FAT-1-3 | 60.022 | 19.902 | Finland | - | 16.0 | 15.6 | Y |
| FI-FAV-1-1 | 60.022 | 19.903 | Finland | - | 14.4 | 14.2 | Y |
| FI-FUT1-2-1 | 60.347 | 27.478 | Finland | - | 16.0 | 16.0 | Y |
| FI-OER-3-3 | 59.789 | 23.174 | Finland | 3.000 | 15.6 | 15.4 | Y |
| FI-SK-17-1 | 59.831 | 23.258 | Finland | 2.500 | 14.4 | 13.8 | Y |
| FI-SK-58-2 | 59.833 | 23.257 | Finland | 0.700 | 14.6 | 14.2 | Y |
| FI-SKW-2-1 | 59.833 | 23.256 | Finland | 1.700 | 15.6 | 15.2 | Y |
| FR-C1-1 | 43.592 | 4.592 | France | - | 18.0 | 17.8 | Y |
| FR-LR5-1 | 43.515 | 3.823 | France | - | 18.0 | 18.0 | Y |
| FR-LR6-1 | 43.452 | 3.808 | France | - | 17.4 | 17.2 | Y |
| FR-LR7-1 | 43.494 | 3.795 | France | - | 18.0 | 17.0 | Y |
| FR-LR8-1 | 43.515 | 3.833 | France | - | 18.0 | 18.0 | Y |
| FR-LR9-1 | 43.567 | 3.970 | France | 0.500 | 17.0 | 16.2 | Y |
| FR-NPM19-1[•] | 46.999 | -2.239 | France | - | 18.0 | 18.2 | Y |
| FR-NPM21-1 | 46.999 | -2.235 | France | - | 18.2 | 17.8 | Y |
| FR-SA-1 | 43.480 | 4.647 | France | - | 18.6 | 17.8 | Y |
| FR-TR-1 | 43.738 | 3.863 | France | - | 17.8 | 17.8 | Y |
| GB-C1-1[•] | 51.734 | -1.336 | England | 0.500 | 13.6 | 13.0 | Y |
| GB-EK1-1 | 55.702 | -2.341 | England | 0.500 | 15.4 | 15.0 | N |
| GB-EK2-6 | 55.698 | -2.343 | England | 0.500 | 15.8 | 15.0 | Y |
| GB-EL75-69 | 51.528 | -0.158 | England | 0.500 | 16.8 | 16.4 | N |
| GB-EP-1 | 52.454 | -1.919 | England | 0.500 | 16.2 | 16.2 | Y |
| GB-FML-1 | 52.531 | -1.956 | England | 0.500 | 16.2 | 16.0 | Y |
| GB-LK1-1 | 55.708 | -2.190 | England | 0.500 | 14.6 | 14.6 | Y |
| GB-S17-7 | 51.620 | -1.386 | England | 0.500 | 16.6 | 16.2 | Y |
| GR-K-1 | 40.704 | 23.144 | Greece | 7.500 | 16.0 | 15.6 | Y |
| HU-AG-03 | 47.515 | 19.081 | Hungary | 0.500 | 15.4 | 15.4 | Y |
| HU-CR3-01 | 47.074 | 19.136 | Hungary | 0.500 | 15.2 | 15.0 | Y |
| HU-K-6 | 46.796 | 19.185 | Hungary | 10.500 | 17.6 | 17.6 | Y |
| HU-KEL-01 | 46.803 | 19.164 | Hungary | - | 16.0 | 15.8 | Y |
| IE-DUB-1 | 53.327 | -6.234 | Ireland | - | 17.4 | 16.8 | Y |
| IL-BM-1 | 30.511 | 34.612 | Israel | - | 16.8 | 16.8 | Y |
| IL-BN-2[•] | 33.143 | 35.774 | Israel | 0.500 | 13.8 | 13.8 | N |
| IL-EY-17 | 30.938 | 35.041 | Israel | 0.500 | 13.6 | 13.2 | Y |
| IL-G-3 | 32.229 | 34.831 | Israel | 1.130 | 14.0 | 14.0 | Y |
| IL-M1-8 | 31.778 | 35.221 | Israel | 0.500 | 17.4 | 17.2 | Y |
| IL-PS-2 | 32.256 | 34.853 | Israel | 1.705 | 17.4 | 17.0 | Y |
| IL-RAM-4 | 33.230 | 35.765 | Israel | 0.500 | 16.6 | 16.6 | Y |
| IL-SH-4 | 31.701 | 34.714 | Israel | - | 16.6 | 16.2 | Y |
| IL-TY-10 | 32.244 | 34.854 | Israel | 4.330 | 14.0 | 14.0 | Y |
| IL-YERU-16 | 30.990 | 34.891 | Israel | - | 16.0 | 15.8 | Y |

| | | | | | | | |
|---|---|---|---|---|---|---|---|
| IT-ISR1-2 | 43.692 | 10.289 | Italy | - | 16.2 | 16.4 | Y |
| IT-MDV-1 | 37.685 | 12.618 | Italy | - | 19.0 | 19.0 | Y |
| IT-PER-2 | 37.519 | 14.307 | Italy | - | 17.4 | 17.4 | Y |
| KG-SK-1 | 41.756 | 75.232 | Kyrgyzstan | - | 17.8 | 17.2 | Y |
| MA-ES-3• | 31.491 | -9.764 | Morocco | - | 18.6 | 18.4 | Y |
| MN-DM1-1 | 45.033 | 100.661 | Mongolia | - | 17.4 | 16.6 | Y |
| NO-AA-1 | 60.051 | 5.074 | Norway | - | 14.4 | 14.2 | Y |
| NO-F-1 | 63.588 | 10.729 | Norway | 0.500 | 15.8 | 15.8 | Y |
| NO-LADE-1 | 63.449 | 10.453 | Norway | 0.500 | 15.4 | 15.0 | Y |
| NO-M3-1 | 59.097 | 11.197 | Norway | - | 16.2 | 16.0 | Y |
| NO-RO-1 | 67.527 | 12.127 | Norway | - | 11.8 | 11.0 | Y |
| NO-V-7 | 67.687 | 12.672 | Norway | - | 14.0 | 13.8 | Y |
| PL-KNP-P4 | 52.323 | 20.731 | Poland | 0.500 | 15.2 | 15.0 | Y |
| PL-W1-1 | 52.211 | 20.997 | Poland | 0.500 | 15.6 | 15.4 | Y |
| PL-W2-1 | 52.230 | 21.028 | Poland | 0.500 | 15.8 | 15.4 | Y |
| RO-DS-3 | 45.147 | 29.678 | Romania | - | 16.0 | 15.6 | Y |
| RU-AL1-11 | 49.968 | 88.692 | Russia | 9.378 | 17.6 | 17.6 | Y |
| RU-AST1-1 | 46.303 | 48.014 | Russia | 0.500 | 16.0 | 15.6 | N |
| RU-AST2-1 | 45.904 | 47.656 | Russia | 0.500 | 15.0 | 15.4 | Y |
| RU-B3-01 | 50.216 | 46.864 | Russia | - | 15.6 | 15.6 | Y |
| RU-B5-1 | 49.986 | 46.693 | Russia | - | 14.4 | 14.4 | Y |
| RU-BAI1-2 | 53.018 | 106.886 | Russia | - | 16.4 | 15.8 | Y |
| RU-BN-BB6 | 50.156 | 43.389 | Russia | - | 15.0 | 15.2 | Y |
| RU-BOL1-1 | 66.426 | 33.837 | Russia | - | 14.6 | 14.8 | Y |
| RU-BOR-1 | 58.075 | 38.199 | Russia | - | 17.4 | 17.2 | Y |
| RU-BP-1 | 58.075 | 38.199 | Russia | - | 16.0 | 15.6 | Y |
| RU-BU2-2 | 51.315 | 108.411 | Russia | 4.730 | 17.6 | 17.4 | Y |
| RU-C10-06 | 50.501 | 47.157 | Russia | - | 13.2 | 13.6 | Y |
| RU-C12-1 | 50.496 | 47.206 | Russia | - | 11.6 | 11.6 | Y |
| RU-C20-1 | 50.510 | 46.519 | Russia | 0.500 | 15.6 | 15.2 | Y |
| RU-HA1-1 | 54.486 | 90.157 | Russia | 4.000 | 17.6 | 17.6 | Y |
| RU-HA2-12 | 54.673 | 90.227 | Russia | 19.000 | 19.4 | 18.8 | Y |
| RU-IRK1-12 | 53.074 | 106.938 | Russia | 2.450 | 16.0 | 16.0 | Y |
| RU-IRK2-14 | 52.989 | 106.721 | Russia | 2.070 | 18.0 | 17.8 | Y |
| RU-IRK3-1 | 53.105 | 107.254 | Russia | 2.657 | 17.2 | 16.4 | Y |
| RU-IRK4-10 | 52.877 | 106.586 | Russia | 21.700 | 19.0 | 18.8 | Y |
| RU-IRK5-72 | 52.835 | 106.580 | Russia | 3.190 | 18.0 | 17.6 | Y |
| RU-IRK9-1 | 53.694 | 102.351 | Russia | - | 17.2 | 17.0 | Y |
| RU-KA2-6 | 45.616 | 45.313 | Russia | 8.210 | 16.4 | 17.0 | Y |
| RU-KOR1-1• | 66.452 | 33.799 | Russia | - | 14.0 | 13.2 | Y |
| RU-KU1-2 | 55.304 | 63.405 | Russia | 1.800 | 14.4 | 13.8 | Y |
| RU-MA10-03 | 46.492 | 42.898 | Russia | 7.346 | 16.0 | 16.0 | Y |
| RU-MA13-3 | 46.444 | 42.809 | Russia | 3.126 | 15.2 | 15.8 | Y |
| RU-MA21-2 | 46.458 | 42.818 | Russia | 12.440 | 16.8 | 16.2 | Y |
| RU-MA3-5 | 46.463 | 42.785 | Russia | 2.400 | 15.4 | 15.2 | Y |
| RU-MA4-3 | 46.439 | 42.745 | Russia | 1.000 | 16.2 | 16.8 | Y |
| RU-MA8-01 | 46.472 | 42.810 | Russia | - | 16.0 | 16.2 | Y |
| RU-NOV1-04 | 55.128 | 77.037 | Russia | - | 17.8 | 17.4 | Y |
| RU-NOV2-01 | 53.754 | 77.894 | Russia | 2.740 | 17.8 | 17.4 | Y |

| | | | | | | | |
|---|---|---|---|---|---|---|---|
| RU-NOV3-2• | 55.032 | 77.466 | Russia | 8.596 | 19.8 | 19.4 | Y |
| RU-NOV4-7 | 53.774 | 78.008 | Russia | - | 18.0 | 17.8 | Y |
| RU-NOV5-17 | 53.746 | 78.053 | Russia | - | 17.8 | 17.8 | Y |
| RU-P1-01 | 54.536 | 39.790 | Russia | 0.500 | 15.2 | 15.2 | Y |
| RU-PIK-2 | 59.995 | 30.473 | Russia | - | 15.8 | 15.6 | Y |
| RU-R2-1 | 56.425 | 37.603 | Russia | 0.500 | 15.2 | 15.0 | Y |
| RU-RM1-2 | 55.763 | 37.582 | Russia | 0.600 | 13.6 | 12.8 | Y |
| RU-RO1-3• | 55.683 | 39.882 | Russia | 0.500 | 13.4 | 13.0 | Y |
| RU-RT1-135 | 45.222 | 36.808 | Russia | - | 16.4 | 16.0 | Y |
| RU-RT21-1-9 | 45.222 | 36.685 | Russia | - | 15.8 | 15.6 | Y |
| RU-SAM2-11 | 52.087 | 50.868 | Russia | - | 12.4 | 12.4 | Y |
| RU-SAM5-1 | 52.923 | 50.317 | Russia | - | 13.4 | 13.0 | Y |
| RU-SPB-35 | 59.811 | 30.133 | Russia | 0.500 | 14.8 | 14.4 | Y |
| RU-SYR1-1 | 53.213 | 48.488 | Russia | 0.500 | 12.0 | 11.8 | Y |
| RU-TU1-1 | 55.624 | 69.209 | Russia | 0.500 | 16.4 | 16.2 | Y |
| RU-TY1-3 | 50.256 | 89.547 | Russia | 14.067 | 15.6 | 15.4 | Y |
| RU-TY3-11 | 51.331 | 93.573 | Russia | 15.630 | 18.0 | 17.8 | N |
| RU-TY4-1 | 50.101 | 95.150 | Russia | - | 17.6 | 17.4 | Y |
| RU-TY5-1 | 52.073 | 93.723 | Russia | 2.188 | 17.8 | 17.6 | Y |
| RU-VOL-36 | 48.530 | 44.487 | Russia | 5.840 | 17.4 | 17.6 | Y |
| RU-YAK1-6 | 61.964 | 129.631 | Russia | 0.500 | 12.2 | 12.2 | Y |
| RU-YAK3-25 | 61.937 | 129.638 | Russia | 0.500 | 12.4 | 12.2 | Y |
| RU-ZB1-1 | 50.413 | 114.725 | Russia | 3.620 | 17.4 | 17.0 | Y |
| RU-ZB2-22 | 50.347 | 114.826 | Russia | 7.720 | 16.4 | 15.4 | Y |
| RU-ZB4-12 | 50.324 | 115.100 | Russia | 5.770 | 17.6 | 17.4 | Y |
| RU-ZB5-1 | 50.318 | 115.305 | Russia | 8.170 | 17.4 | 17.4 | Y |
| RU-ZB6-03 | 50.226 | 115.642 | Russia | 1.906 | 17.6 | 17.6 | Y |
| SE-BY-J6 | 55.675 | 13.545 | Sweden | - | 16.2 | 15.8 | Y |
| SE-G1-9 | 60.422 | 18.510 | Sweden | 1.000 | 11.4 | 10.8 | Y |
| SE-G2-8 | 60.432 | 18.522 | Sweden | 1.000 | 14.8 | 14.4 | Y |
| SE-G3-7 | 60.423 | 18.557 | Sweden | 1.000 | 14.8 | 14.4 | N |
| SE-G4-20 | 60.415 | 18.569 | Sweden | 1.000 | 14.6 | 14.0 | Y |
| SE-GN1-3d3 | 60.497 | 18.431 | Sweden | 1.500 | 15.4 | 15.4 | Y |
| SE-GN2-3A10 | 60.497 | 18.432 | Sweden | 1.500 | 14.8 | 14.6 | Y |
| SE-H1-1 | 58.342 | 11.218 | Sweden | 2.100 | 15.8 | 16.0 | N |
| TN-RA-2 | 36.960 | 10.224 | Tunisia | - | 18.2 | 18.2 | Y |
| TR-EG-1 | 39.824 | 32.831 | Turkey | 3.800 | 16.6 | 16.2 | Y |
| UA-KR1-7 | 45.094 | 36.311 | Ukraine | - | 16.4 | 16.4 | Y |
| US-D-1 | 38.530 | -121.611 | USA | - | 16.0 | 15.8 | Y |
| US-SP15-1 | 44.333 | -68.063 | USA | - | 14.2 | 14.2 | Y |
| US-SP163-1 | 44.334 | -68.064 | USA | - | 14.0 | 14.0 | Y |
| US-SP2-1 | 44.333 | -68.063 | USA | - | 14.0 | 14.0 | Y |
| US-SP221-1 | 44.334 | -68.065 | USA | - | 14.0 | 13.8 | Y |
| US-SP6-10 | 44.334 | -68.064 | USA | - | 14.0 | 14.0 | Y |
| US-SP7-1 | 44.332 | -68.061 | USA | - | 14.0 | 13.2 | Y |

**Table S2. Significant outliers SNPs for salinity tolerance in *D. magna*.** The significant threshold for the genome-wide association study (gwas) was based on a Bonferroni-adjusted *p*-value=$3.4 \times 10^{-9}$. Each SNP is identified by the chromosome (CHR), contig, position in the contig (Pos_contig) and positioSn in the entire genome (Pos_genome) where it occurs. The *p*-value is presented in the form of –Log10(*p*).

| Chromosome | Contig | Pos_contig | Pos_genome | -Log10(*p*) | Candidate gene |
|---|---|---|---|---|---|
| 7 | 000003F | 2931307 | 84915987 | 9.263 | 3' UTR uncharacterized |
| 7 | 000003F | 2931312 | 84915982 | 9.249 | 3' UTR uncharacterized |
| 7 | 000003F | 2931372 | 84915922 | 8.835 | 3' UTR uncharacterized |
| 7 | 000003F | 2931486 | 84915808 | 8.632 | 3' UTR uncharacterized |
| 7 | 000003F | 2932601 | 84914693 | 8.822 | 5' UTR *Vav3 GEF* |
| 7 | 000003F | 2932686 | 84914608 | 9.071 | 5' UTR *Vav3 GEF* |
| 7 | 000003F | 2932763 | 84914531 | 9.024 | 5' UTR *Vav3 GEF* |
| 7 | 000003F | 2932770 | 84914524 | 9.658 | 5' UTR *Vav3 GEF* |
| 7 | 000003F | 2932785 | 84914509 | 10.139 | *TPS* |
| 7 | 000003F | 2932870 | 84914424 | 9.829 | TPS |
| 7 | 000003F | 2932920 | 84914374 | 9.683 | TPS |
| 7 | 000003F | 2932925 | 84914369 | 9.627 | TPS |
| 7 | 000003F | 2932962 | 84914332 | 8.502 | TPS |
| 7 | 000003F | 2933798 | 84913496 | 8.957 | TPS |
| 7 | 000003F | 2933800 | 84913494 | 8.781 | TPS |
| 7 | 000003F | 2933912 | 84913382 | 8.570 | TPS |
| 7 | 000003F | 2933949 | 84913345 | 8.666 | TPS |
| 7 | 000003F | 2933959 | 84913335 | 8.500 | TPS |
| 7 | 000003F | 2934396 | 84912898 | 8.558 | TPS |
| 7 | 000003F | 2934397 | 84912897 | 8.697 | TPS |
| 7 | 000003F | 2934407 | 84912887 | 8.697 | TPS |
| 7 | 000003F | 2934409 | 84912885 | 8.697 | TPS |
| 7 | 000003F | 2934451 | 84912843 | 9.793 | TPS |
| 7 | 000003F | 2934705 | 84912589 | 10.169 | TPS |
| 7 | 000003F | 2934735 | 84912559 | 10.103 | TPS |
| 7 | 000003F | 2934830 | 84912464 | 11.648 | TPS |
| 7 | 000003F | 2934983 | 84912311 | 9.581 | TPS |

**Table S3. The sense oligonucleotide sequences for sgRNA synthesis.** T7 promoter, target sequence, and the first 20 bp of the Cas9 binding scaffold sequence are indicated with bold letters, underline, and italic letters respectively.

| sgRNA Target | Sense oligonucleotide |
|---|---|
| sgRNA#1 | 5′-**GAAATTAATACGACTCACTATA** **GC**GATAAGTCTTTCCTGCGA*GTTTTAGAGCTAGAAA*-3′ |
| sgRNA#2 | 5′-**GAAATTAATACGACTCACTATA** **GA**CTGTGGCGGGCGATGGGT*GTTTTAGAGCTAGAA*-3′ |

**Table S4.** Primers for genotyping the mutant genotype, including information on target region, amplification direction and primer sequence.

| Target region | Direction | Sequence (5′ → 3′) |
|---|---|---|
| sgRNA#1 | Forward | GTGTAAGGGTAGATGGTGGTAGG |
| | Reverse | CCAGTTACATCGTCACGTTTG |
| sgRNA#2 | Forward | AGGTCTCGTGACTGCCGTTG |
| | Reverse | GTTTCAGCAGACCAGCCGTG |

**Table S5. Protein sequences of the TPS gene in NIES (control) and TPS mutant genotypes**. Residues in underlined and italics indicate the G6P entry site and inappropriate amino acid insertions, respectively. Codon stops are represented by an asterisk.

| Genotype | TPS protein sequence |
|---|---|
| NIES<br>(control 1 and 2) | MALNVHQESKSEVEWNPSQERLIVVSNRLPFVLKRDDVTGKLVRKSSAGGLVTAVAPVVVDCGGR<br>WVGWTGDCSFGVEDEIPEASEEDTTPTAGLLKHAQVVPVNLQQQFDSYYNGCCNGTLWPLFHS<br>MSDRAVYSTEFWQDYIQVNQLFADKTLEAIELTSAEGPGIVSLVWIHDYHLTLLPSILRQALSENNC<br>NARLGFFLHIPFPPWDIFRLLPWDDEILLGLLFCDMIGFHIEDYCINFLDCCHRRLGCRIDRKNLLVEY<br>GNRIIKIAALPIGIPFQRFQKMAEEAPRLIKEKRQKIILGVDRLDYTKGLVHRLKAFERFLEKHPEHIEE<br>VIFIQIAVPSRTDVKEYQDLKEEIDQVIGRINGRFSTANWSPIRYIYGCVSQSELAAYYRDAFIAMVTP<br>LRDGMNLVAKEFVACQIEEPGVLILSPFAGAGERMQEALLINPYKIDDVAELICSALEMSRDEREVR<br>MKNLRRREKMNNVDAWAQSFLKTIRAITSDQGSDSKMGSSLMSSAMDDYDEYLSKYVGNAEKL<br>ALLLDYDGTLAPLAPHPDLAILPNETGKILHRLSNCPDVYISIISNRSVENVKKMVGIENITYAGKNGL<br>EILHPDFTQFIYPLPVEYEDKVRSLLRQLQEEVCNDGAWVEHKGLLLTYHYSETVMHLRTGLIKRAR<br>ELIEGAGLVCTTAYCALEAKPPSQWNKGQAALYILRTAFGVDWEERIRIVYAGDDFTDEAAIKALKG<br>LAATFRVTSTLAVKTAADRRLSNNDSVLRMLKWIESHMTMRKPRLSSRSSCTVLE1LDEDQDPST |
| TPS mutant A –<br>Allele 1 | MALNVHQESKSEVEWNPSQERLIVVSNRLPFVLKRDDVTGKLVRKSSAGGLVTAVAPVVVDCGGR<br>SWMDG* |
| TPS mutant A –<br>Allele 2 | MALNVHQESKSEVEWNPSQERLIVVSNRLPFVLKRDDVTGKLVRKSSAGGLVTAVAPVVVGVEDE<br>IPEASEEDTTPTAGLLKHAQVVPVNLQQQFDSYYNGCCNGTLWPLFHSMSDRAVYSTEFWQDYI<br>QVNQLFADKTLEAIELTSAEGPGIVSLVWIHDYHLTLLPSILRQALSENNCNARLGFFLHIPFPPWDIF<br>RLLPWDDEILLGLLFCDMIGFHIEDYCINFLDCCHRRLGCRIDRKNLLVEYGNRIIKIAALPIGIPFQRF<br>QKMAEEAPRLIKEKRQKIILGVDRLDYTKGLVHRLKAFERFLEKHPEHIEEVIFIQIAVPSRTDVKEYQ<br>DLKEEIDQVIGRINGRFSTANWSPIRYIYGCVSQSELAAYYRDAFIAMVTPLRDGMNLVAKEFVACQ<br>IEEPGVLILSPFAGAGERMQEALLINPYKIDDVAELICSALEMSRDEREVRMKNLRRREKMNNVDA<br>WAQSFLKTIRAITSDQGSDSKMGSSLMSSAMDDYDEYLSKYVGNAEKLALLLDYDGTLAPLAPHPD<br>LAILPNETGKILHRLSNCPDVYISIISNRSVENVKKMVGIENITYAGKNGLEILHPDFTQFIYPLPVEYE<br>DKVRSLLRQLQEEVCNDGAWVEHKGLLLTYHYSETVMHLRTGLIKRARELIEGAGLVCTTAYCALE<br>AKPPSQWNKGQAALYILRTAFGVDWEERIRIVYAGDDFTDEAAIKALKGLAATFRVTSTLAVKTAA<br>DRRLSNNDSVLRMLKWIESHMTMRKPRLSSRSSCTVLE1LDEDQDPST |
| TPS mutant B –<br>Allele 1 | MALNVHQESKSEVEWNPSQERLIVVSNRLPFVLKRDDVTGKLVRKSSARV* |
| TPS mutant B –<br>Allele 2 | MALNVHQESKSEVEWNPSQERLIVVSNRLPFVLKRDDVTGKLVRKSSAGGLVTAVAPVVVGCGGR<br>*CLSQPPWSILV*GWTGDCSFGVEDEIPEASEEDTTPTAGLLKHAQVVPVNLQQQFDSYYNGCCNGT<br>LWPLFHSMSDRAVYSTEFWQDYIQVNQLFADKTLEAIELTSAEGPGIVSLVWIHDYHLTLLPSILRQ<br>ALSENNCNARLGFFLHIPFPPWDIFRLLPWDDEILLGLLFCDMIGFHIEDYCINFLDCCHRRLGCRID<br>RKNLLVEYGNRIIKIAALPIGIPFQRFQKMAEEAPRLIKEKRQKIILGVDRLDYTKGLVHRLKAFERFLE<br>KHPEHIEEVIFIQIAVPSRTDVKEYQDLKEEIDQVIGRINGRFSTANWSPIRYIYGCVSQSELAAYYRD<br>AFIAMVTPLRDGMNLVAKEFVACQIEEPGVLILSPFAGAGERMQEALLINPYKIDDVAELICSALEM<br>SRDEREVRMKNLRRREKMNNVDAWAQSFLKTIRAITSDQGSDSKMGSSLMSSAMDDYDEYLSKY<br>VGNAEKLALLLDYDGTLAPLAPHPDLAILPNETGKILHRLSNCPDVYISIISNRSVENVKKMVGIENIT<br>YAGKNGLEILHPDFTQFIYPLPVEYEDKVRSLLRQLQEEVCNDGAWVEHKGLLLTYHYSETVMHLR<br>TGLIKRARELIEGAGLVCTTAYCALEAKPPSQWNKGQAALYILRTAFGVDWEERIRIVYAGDDFTDE<br>AAIKALKGLAATFRVTSTLAVKTAADRRLSNNDSVLRMLKWIESHMTMRKPRLSSRSSCTVLE1LDE<br>DQDPST |

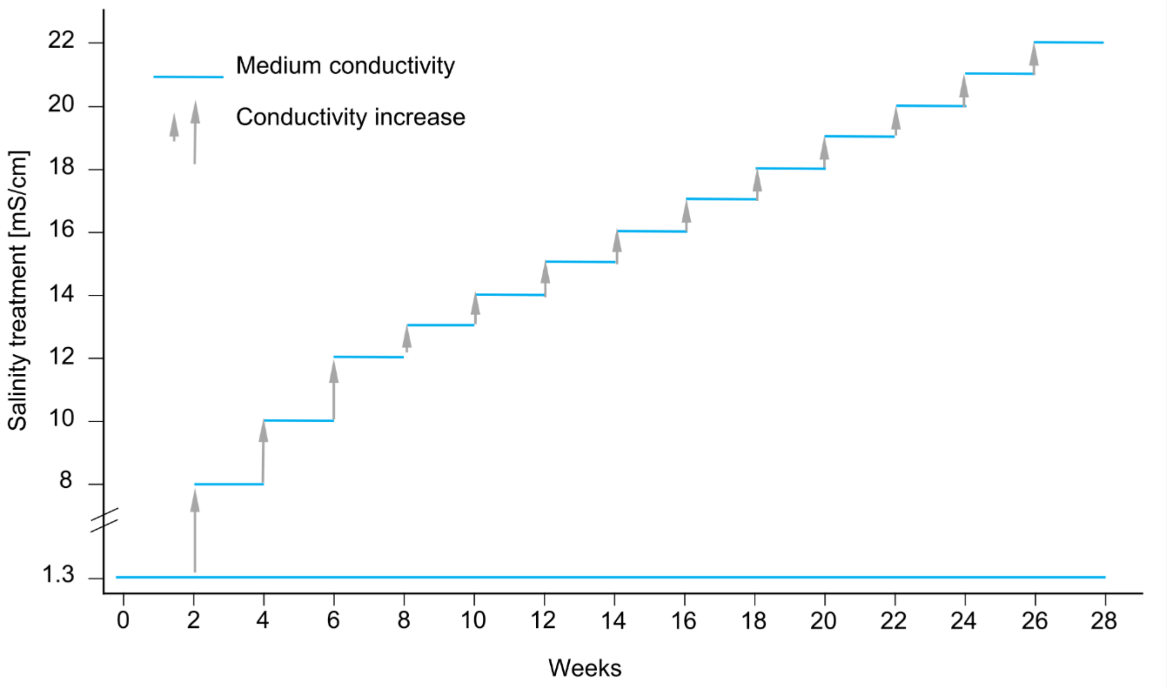


**Figure S1. Experimental design for the** determination of salinity tolerance in *D. magna*. Animals were taken from stock cultures and cultured in 360-mL jars filled with 350 mL *Daphnia* medium (ADaM, 1.3 mS/cm, 20 °C). Every two weeks, the population of a replicate was transferred to medium with higher salinity, and population survival and newborn were recorded. If at least one animal of a replicate survived the 14-day assay period in a given salinity treatment, juveniles from that replicate were tested at the next higher salinity level. If all animals died, we assumed they exceeded their upper salinity tolerance. As salinity increased, fewer and fewer replicate lines survived. Control animals were kept in ADaM. All control lines survived.

**A**

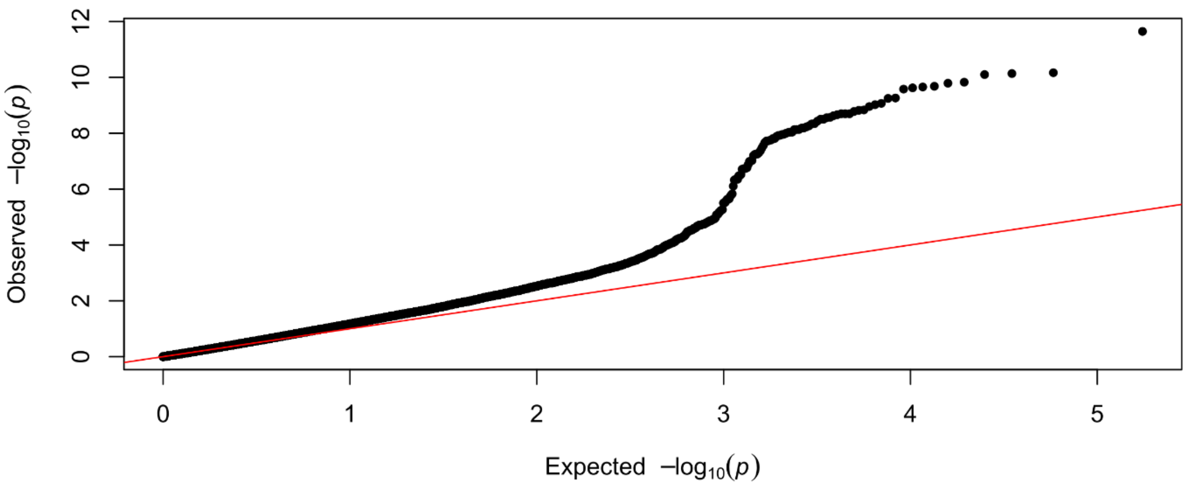


**B**

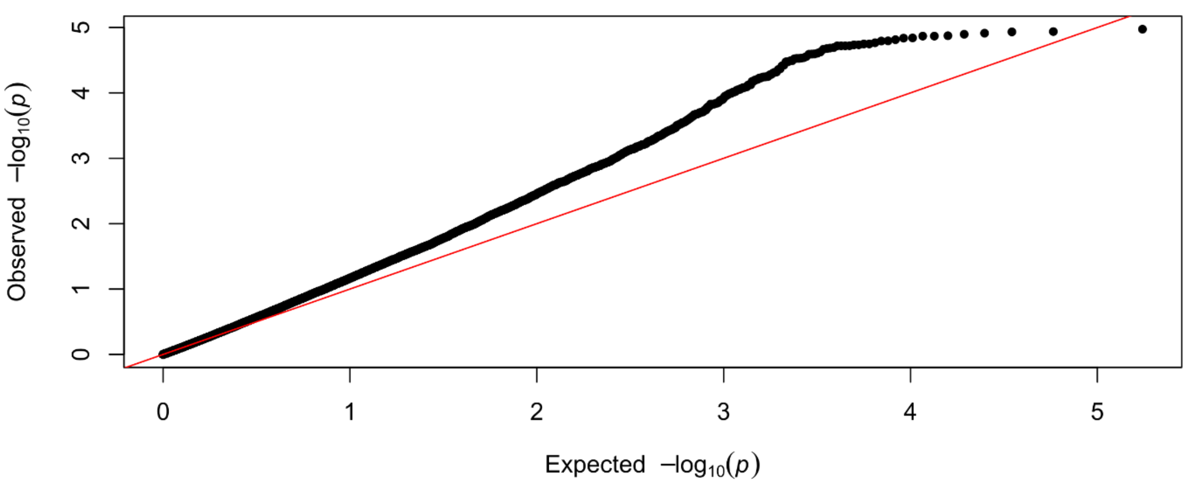


**Figure S2.** Quantile-quantile (QQ) plots of the genome-wide association study for salinity tolerance. *P*-value distributions for the whole contig 000003F on chromosome 7 (A) and the same contig excluding the peak region (i.e., *p*-value=5.0x10$^{-5}$), compared to the expected *p*-values under a uniform distribution (red). After the exclusion of the significantly associated region, the observed *p*-values relate more to the expected *p*-values.

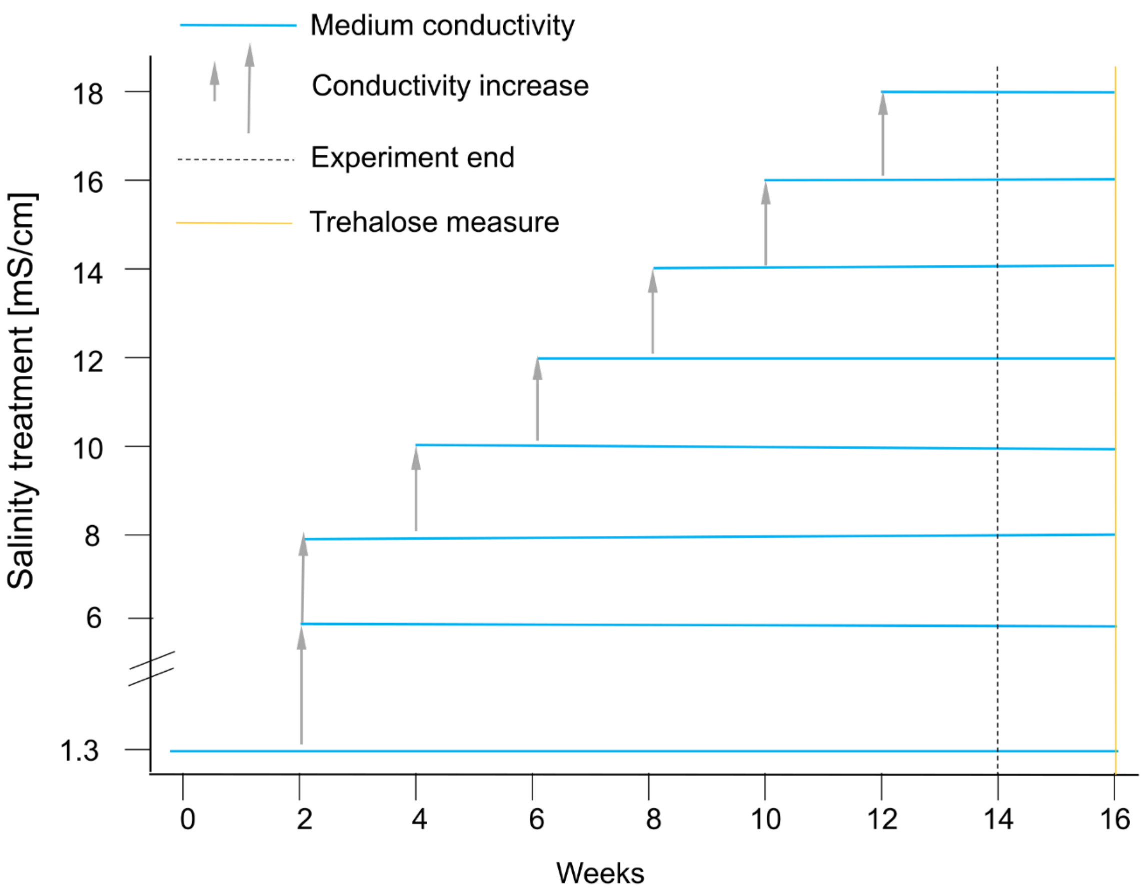


**Figure S3. Experimental design for measuring survival, trehalose and protein concentration**. Animals were taken from stock cultures (at 20 °C) and cultured in 360-mL jars filled with 350 mL *Daphnia* medium (ADaM, 1.3 mS/cm, 20 °C). At week 2, replicate populations with a salinity level of 6 and 8 mS/cm were produced. Every two weeks, replicate populations in the highest salinity treatment were divided and one group was transferred to medium with 2 mS/cm of salinity. Population survival was monitored. If at least one animal of a replicate survived the 14-day assay in the highest salinity, juveniles from that replicate were tested at the next higher salinity level. If all animals died, we assumed that they had exceeded their upper salinity tolerance. Four weeks after all replicates had reached their upper salinity tolerance, we quantified trehalose and protein concentrations in animals from all replicates in salinity treatments (1.3, 8, 10, 12, 14, 16 and 18 mS/cm) (yellow line in the figure). As salinity increased, fewer and fewer replicates survived to be tested.

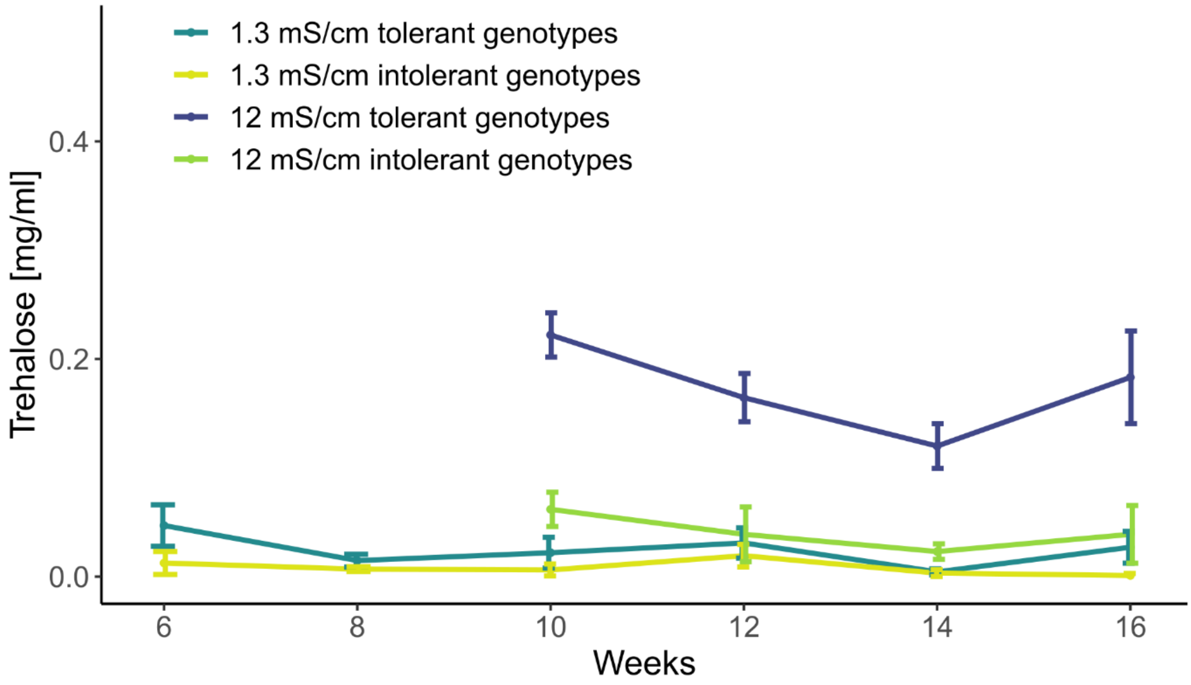


**Figure S4. Weekly variation in trehalose concentration.** Measurements were taken every two weeks for salinity tolerant and intolerant *D. magna* genotypes at salinity treatments of 1.3 and 12. There is no significant variation in trehalose concentration among weeks (Df=1, F-value=1.60, *p*-value=0.22). Artificial Daphnia Medium (ADaM) has a salinity of 1.3 mS/cm.

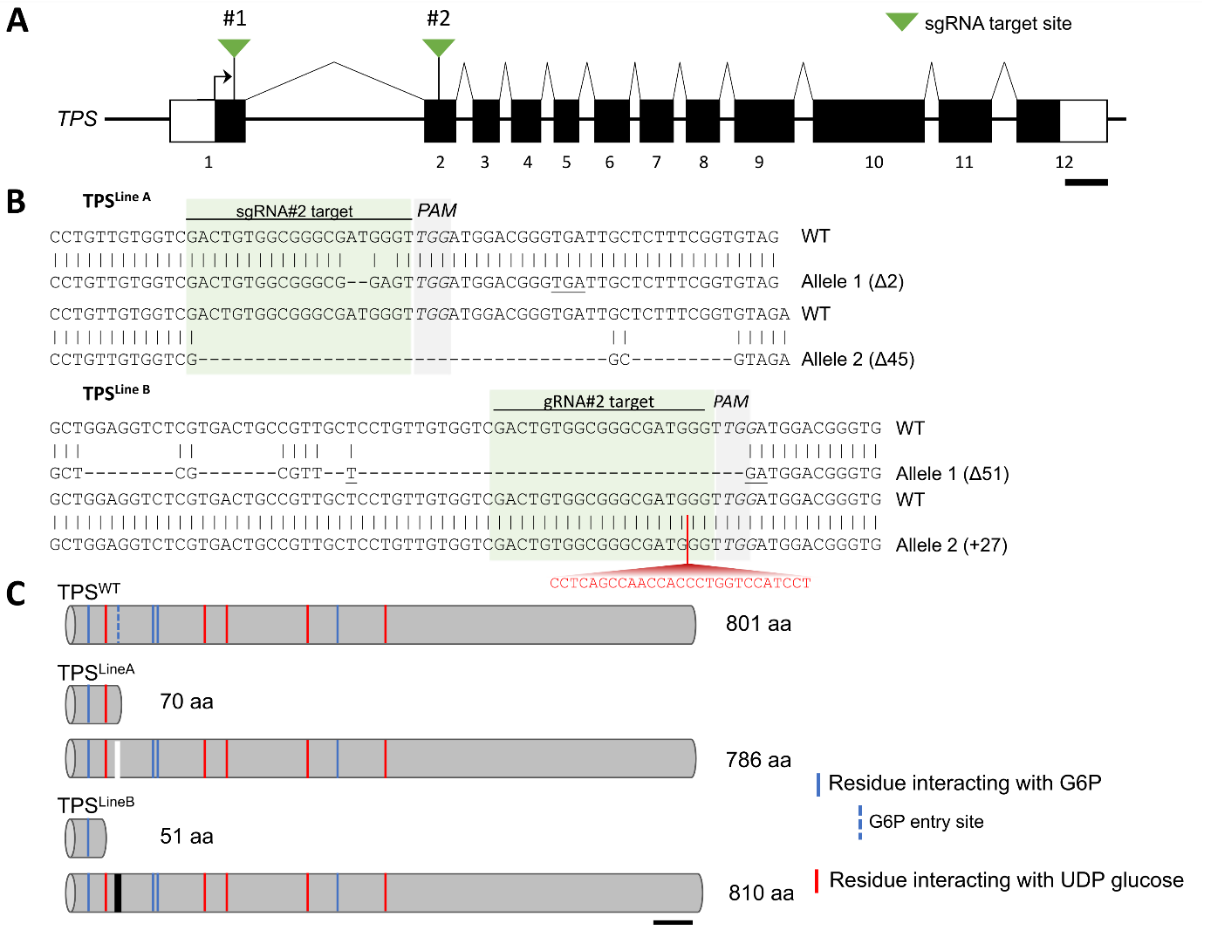


**Figure S5. Generation of TPS mutant by CRISPR/Cas9 system**. (A) Genomic structure of the TPS gene and sgRNAs target sites. Exons are indicated as black, the transcription initiation site as an arrow, and sgRNA target sites as green triangles. Data are derived from *D. magna* (SK strain) genome assembly (NCBI accession number XM_ 032933135.1). (B) The sequence of sgRNA#2 target site and mutated regions of two isolated TPS mutant genotypes: TPS mutant A and B. The 20 bp sgRNA spacer is indicated in green, PAM in grey, and mutated or deleted bases in red and hyphens, respectively. Locations of premature stop codons caused by indel mutation are underlined. (C) Predicted translated protein from TPS mutant alleles. Blue and red lines indicate residue interacting with substrate glucose-6-phosphate (G6P) and UDP glucose, respectively. The blue-dashed line indicates the G6P entry site. TPS mutant A produces two types of proteins: 70 amino acids (aa) which lack most of the catalytic residues and 786 aa with an in-frame deletion in the entire G6P entry site (white bar). TPS mutant B alleles translated into proteins with 51 aa and 810 aa. Similarly, the short protein lacks all but the first catalytic residue, while the long one has in-frame inappropriate amino acids inserted in the G6P entry site (black bar). Scale bar in panels A and C are 200 nt are 50 aa, respectively.
